# Brief and diverse excitotoxic insults cause an increase in neuronal nuclear membrane permeability in the neonatal brain

**DOI:** 10.1101/2023.08.22.554167

**Authors:** P. Suryavanshi, R. Langton, K. Fairhead, J. Glykys

## Abstract

Neuronal swelling after excitotoxic insults is implicated in neuronal injury and death in the developing brain, yet mitigating brain edema with osmotic and surgical interventions yields poor clinical outcomes. Importantly, neuronal swelling and its downstream consequences during early brain development remain poorly investigated. Using multiphoton Ca^2+^ imaging *in vivo* (P12-17) and in acute brain slices (P8-12), we explored Ca^2+^-dependent downstream effects after neuronal cytotoxic edema. We observed the translocation of cytosolic GCaMP6s into the nucleus of a subpopulation of neurons minutes after various excitotoxic insults. We used automated morphology-detection algorithms for neuronal segmentation and quantified the nuclear translocation of GCaMP6s as the ratio of nuclear and cytosolic intensity (N/C ratio). Elevated neuronal N/C ratios were correlated to higher Ca^2+^ loads and could occur independently of neuronal swelling. Electron microscopy revealed that the nuclear translocation was associated with increased nuclear pore size. Inhibiting calpains prevented elevated N/C ratios and neuronal swelling. Thus, our results indicate altered nuclear transport in a subpopulation of neurons shortly after injury in the developing brain, which can be used as an early biomarker of acute neuronal injury.

## Introduction

Multiple brain insults, including stroke, hypoxia, prolonged seizures, and traumatic brain injury, cause brain swelling, which is associated with severe morbidity or death (Battey et al., 2014; Cooper et al., 2011; Stokum et al., 2016; Unterberg et al., 2004). Cytotoxic edema during early brain development often leads to long-lasting motor and cognitive deficits in survivors (Gonzalez and Miller, 2006; Low et al., 1988). Osmotic and surgical interventions employed to mitigate cellular edema continue to yield poor outcomes (Hutchinson et al., 2016; Kolias et al., 2022), suggesting that mechanisms independent of brain swelling may contribute to cell injury and death (Cooper et al., 2011; Ferrari et al., 2010; Hutchinson et al., 2016). Notably, there is a lack of therapeutic strategies targeting neonates and children, and current treatments have been extrapolated from adult studies. Thus, there is a critical need to understand the downstream consequences of neuronal swelling leading to poor outcomes, especially in pediatric patients.

In addition to swelling, excitotoxic insults can induce intracellular Ca^2+^ elevation in neurons (Berridge et al., 2003). Rapid neuronal Ca^2+^ influx following neuronal activity and excitotoxic Ca^2+^ overload can be visualized using genetically encoded fluorescent Ca^2+^ indicators (GECIs) like GCaMPs (Akerboom et al., 2012; Chen et al., 2012). Without nuclear localization signals, macromolecules like GCaMPs (~70 kD) are restricted to the cytosol, as only smaller proteins can permeate the nucleus through a passive diffusion (Miao and Schulten, 2009; Wente and Rout, 2010). Therefore, nuclear accumulation of typically cytosolic GCaMPs, which is sometimes observed (Yang et al., 2018), could suggest modified nuclear membrane permeability (Strasser et al., 2012; Yamashita et al., 2017). Neurons with GCaMP-filled nuclei also show an altered physiology (Resendez et al., 2016; Tian et al., 2009). However, previous research in cultured neuronal or non-neuronal cells indicates that nuclear membrane breakdown and consequent nuclear accumulation of macromolecules is a slow process that typically requires prolonged exposure to supraphysiological insults (Bano et al., 2010; Ferrando-May et al., 2001; Strasser et al., 2012).

In this study, we demonstrate *ex-* and *in vivo,* a rapid calpain-dependent nuclear translocation of GCaMP6s following brief and physiologically relevant excitotoxic insults recapitulating various neurological conditions (some associated with cytotoxic edema) during early brain development. The nuclear translocation of GCaMP6s was associated with enlarged nuclear pores in neocortical neurons. Our results demonstrate that the abnormal translocation of GCaMP6s to the nucleus following excitotoxic insults occurs in minutes and is independent of neuronal swelling, indicating dysregulation of protein localization in injured neurons. Our results shed light on some reasons for the lack of good clinical outcomes even after mitigating brain swelling. Additionally, since the expression of biomarkers associated with neuronal death and injury is typically delayed (hours to days), we also propose that this abnormal nuclear translocation can be a valuable tool for the early detection of neuronal injury.

## Results

### Brief NMDA exposure induces long-term swelling and Ca^2+^ overload in neocortical neurons during early brain development

Various brain insults during early brain development involve NMDA receptor-mediated excitotoxicity. Using multiphoton fluorescent imaging, we evaluated excitotoxic consequences of brief NMDAR activation on neuronal swelling and altered Ca^2+^ dynamics in mouse neocortical neurons (layer IV/V, brain slices) during early brain development (Thy1-GCaMP6s, P8-12). A brief application of NMDA (30 µM for 10 min) induced hyperexcitability, prolonged neuronal Ca^2+^ elevation (**Fig. S1A, B**) and caused varicose swelling or beading of the dendrites (Thy1-YFP, **Fig. S1C**).

To simultaneously image neuronal edema and Ca^2+^ activity, we expressed a neuron-specific stable fluorophore (mRuby2) and a Ca^2+^ sensor (GCaMP6s) (Rose et al., 2016), using intraventricular AAV injections in newborn pups (P0-2; **Fig. 1A**) (Glascock et al., 2011). Neuronal fluorophore expression was evident at 7 days post-injection. Neuronal areas were measured using an automated convolutional neuronal network algorithm (ANMAF; **Fig. 1A**), which detects and measures neuronal size while avoiding human bias in the area determination (Tong et al., 2021). NMDA application triggered synchronized Ca^2+^ transients in neurons followed by a persistently high Ca^2+^ after 40 min of washout in a subset of neurons (**Fig. 1B and C, Video S1**). NMDA perfusion also caused a persistent increase in neuronal area (**Fig. 1D, Fig. S2A-D**). Ca^2+^ elevation had a weak but significant correlation with swelling during but not after NMDA perfusion (**Fig. S2E**). Notably, there was no significant difference when either GCaMP6s or mRuby2 was used to measure changes in neuronal size (**Fig. 1D**).

**Fig. 1:**
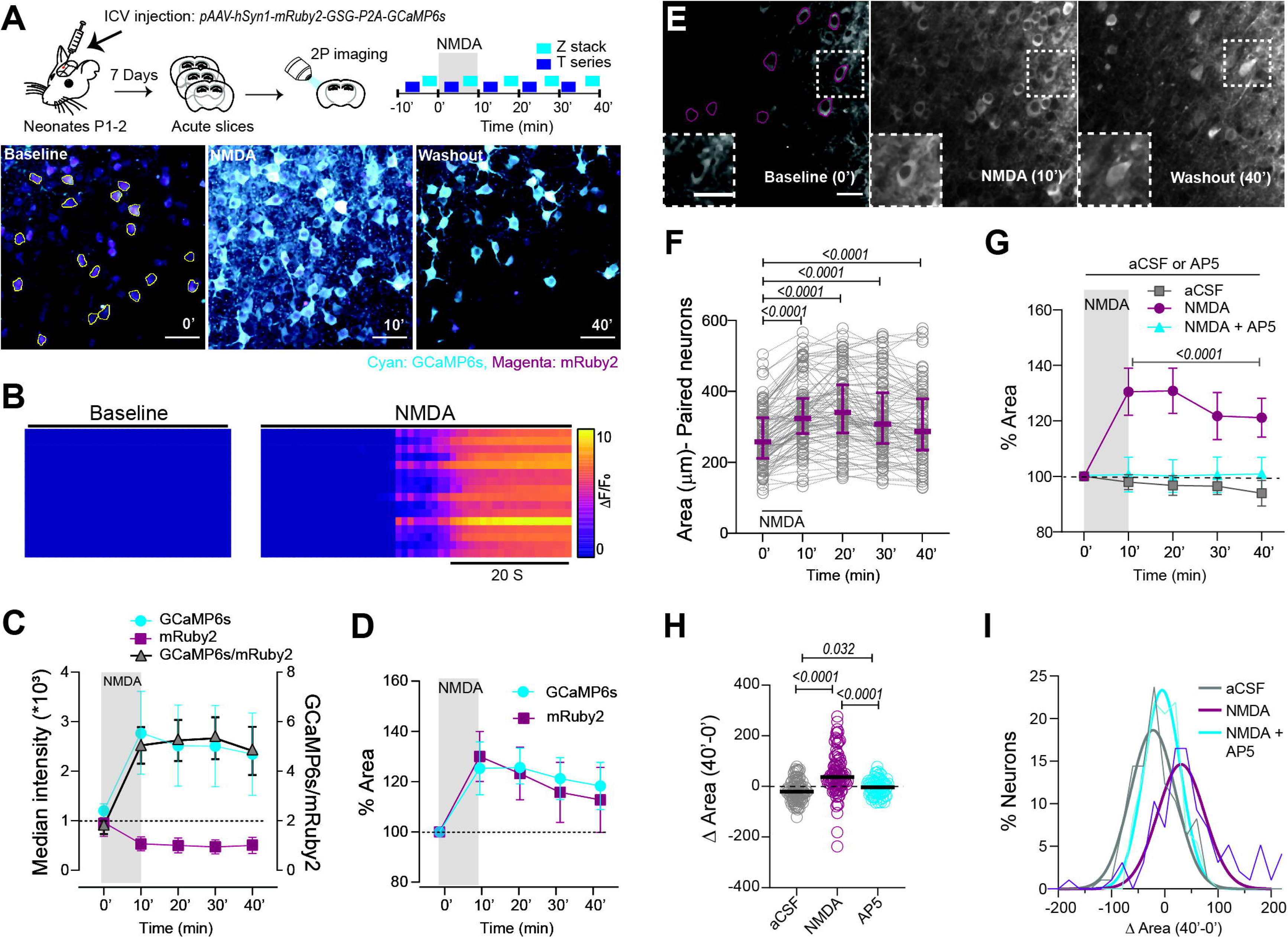
Brief NMDA perfusion induces long-term neuronal swelling and increases neuronal Ca^2+^ in neocortical neurons. **(A)** *Top:* neuron-specific viral delivery of a stable fluorophore (mRuby2) and a Ca^2+^ sensor (GCaMP6s) along with the experimental design describing the acquisitions of z-stacks and single-plane t-series. *Bottom:* representative dual-color images of mRuby2 and GCaMP6s expressing cortical neurons before (0’), during (10’), and after (40’) NMDA application, with ANMAF-generated ROIs overlayed on the baseline image. **(B)** Synchronized Ca^2+^ activity (ΔF/F_0_) heatmap of neurons during NMDA application (n=27 neurons from 1 slice, **Video S1)**. **(C)** Long-lasting increase in GCaMP6s signals along with **(D)** persistent elevation of the neuronal area over 40’ after NMDA treatment, detected using both GCaMP6s and mRuby2 *(n [mice: slices: neurons]=2:3:54).* **(E)** Representative images of GCaMP6s-expressing neocortical neurons at different time points (0’, 10’, and 40’). *Insets*: A single neuron at higher magnification shows persistent elevation in Ca^2+^. **(F)** Matched comparisons of neuronal areas between multiple time points show persistent swelling in most neurons. RM one-way ANOVA with Dunnett’s post-test, F(2.8, 261)=49.6, p<0.0001. **(G)** Significant and persistent increase in the neuronal areas after NMDA application, prevented by AP5 treatment. Two-way ANOVA with Tukey’s post-test, Interaction: F (8, 1020) =12.1, p<0.0001, Treatment: F (2, 1020) =165.3, p<0.0001, Time: F (4, 1020) =9.9, p<0.0001; n = 5:8:97 (aCSF), n = 6:11:97 (NMDA), n = 3:5:73 (AP5). **(H)** Increase in areas of paired neurons at 40’ after NMDA application (Δ Area, 40’ minus 0’), rescued partially by AP5. Kruskal-Wallis test with Dunn’s post-test, p<0.0001, ‘n’ same as **Fig. 1G**.**(I)** Distribution of total neuronal Δ Area at 40’ (n: aCSF=579 neurons, NMDA=315, AP5=123). Data represented as mean ± 95% CI (parametric data) or median ± IQR (non-parametric data). Scale bar: 50 µm. See **also Fig. S1 and S2**.

As neuronal swelling measurements were similar between mRuby2 and GCaMP6s, we turned to transgenic mice expressing GCaMP6s in a subset of neocortical neurons (Thy1-GCaMP6s) for the rest of our experiments (**Fig. 1E**). We confirmed that brief NMDA application in acute neocortical slices from Thy1-GCaMP6s mice (P8-12) caused long-term neuronal swelling in a subset of neurons (**Fig. 1F and G**). In contrast, other neurons showed resistance to NMDA-induced excitotoxicity or a return to baseline (**Fig. 1F, H-I, Supplementary Fig. 3C**). NMDA-induced neuronal swelling and [Ca^2+^]_i_ elevation was blocked by the NMDA receptor antagonist AP5. Swelling and elevated [Ca^2+^]_i_ was also absent during prolonged perfusion with artificial cerebrospinal fluid (aCSF, **Fig. 1G-I, Supplementary Fig. 3A-C**), indicating that the persistent swelling and Ca^2+^ elevation are caused by transient NMDA receptor activation. With aCSF, but not AP5, there was a slight decrease in the neuronal area over time, possibly due to the aCSF osmolarity (300 mOsm). In summary, these results show that excitotoxic neuronal swelling and Ca^2+^ elevation occur concurrently, and that neuronal area can be measured with GCaMP6s.

### Excitotoxic injury to neurons causes nuclear translocation of cytosolic GCaMP6s in the developing neocortex

Although GCaMPs are widely used GECIs due to their rapid response kinetics and good signal-to-noise ratio (Akerboom et al., 2012), few reports show an abnormal and detrimental nuclear accumulation of GCaMP in a few neurons (Tian et al., 2009; Yang et al., 2018). To avoid these “injured” neurons with GCaMP-filled nuclei in the superficial tissue of our acute slices (~0 to 40 µm from the surface; **Fig. S1D**), we restricted our imaging to healthy neurons (cytosolic GCaMP6s expression) located at 80-200 µm below the slice surface. Interestingly, we observed gradual GCaMP6s translocation to the nucleus in a subset of neurons following NMDA application, even at optimal imaging depths (**Fig. 2A**). To quantify the changes in nuclear GCaMP6s intensity, we normalized the nuclear to the cytosolic GCaMP6s fluorescence (N/C ratio), using compartmentalized ROIs (**Fig. 2B**). The N/C ratio was unchanged in neurons under prolonged aCSF perfusion yet was significantly increased after NMDA perfusion and prevented by AP5 (**Fig. 2C-E**). The elevation in the N/C ratio began shortly after NMDA treatment and persisted for the duration of the experiment (~30 minutes, **Fig. S4A-D**). The N/C ratio did not correlate to changes in neuronal size (at 10 and 40 minutes, **Fig. S2F**) but correlated to an increase in Ca^2+^ load at 40 minutes (ΔN/C ratio vs. ΔGreen/Red ratio, **Fig. 2F**), suggesting a possible Ca^2+^-mediated mechanism underlying the NMDA-induced increase in the N/C ratio. Interestingly, the N/C ratio increased in some neurons that shrank during aCSF perfusion alone (at 40 minutes, **Fig. S3D**), further negating a relationship between the degree of neuronal swelling and elevated N/C ratio.

**Fig. 2:**
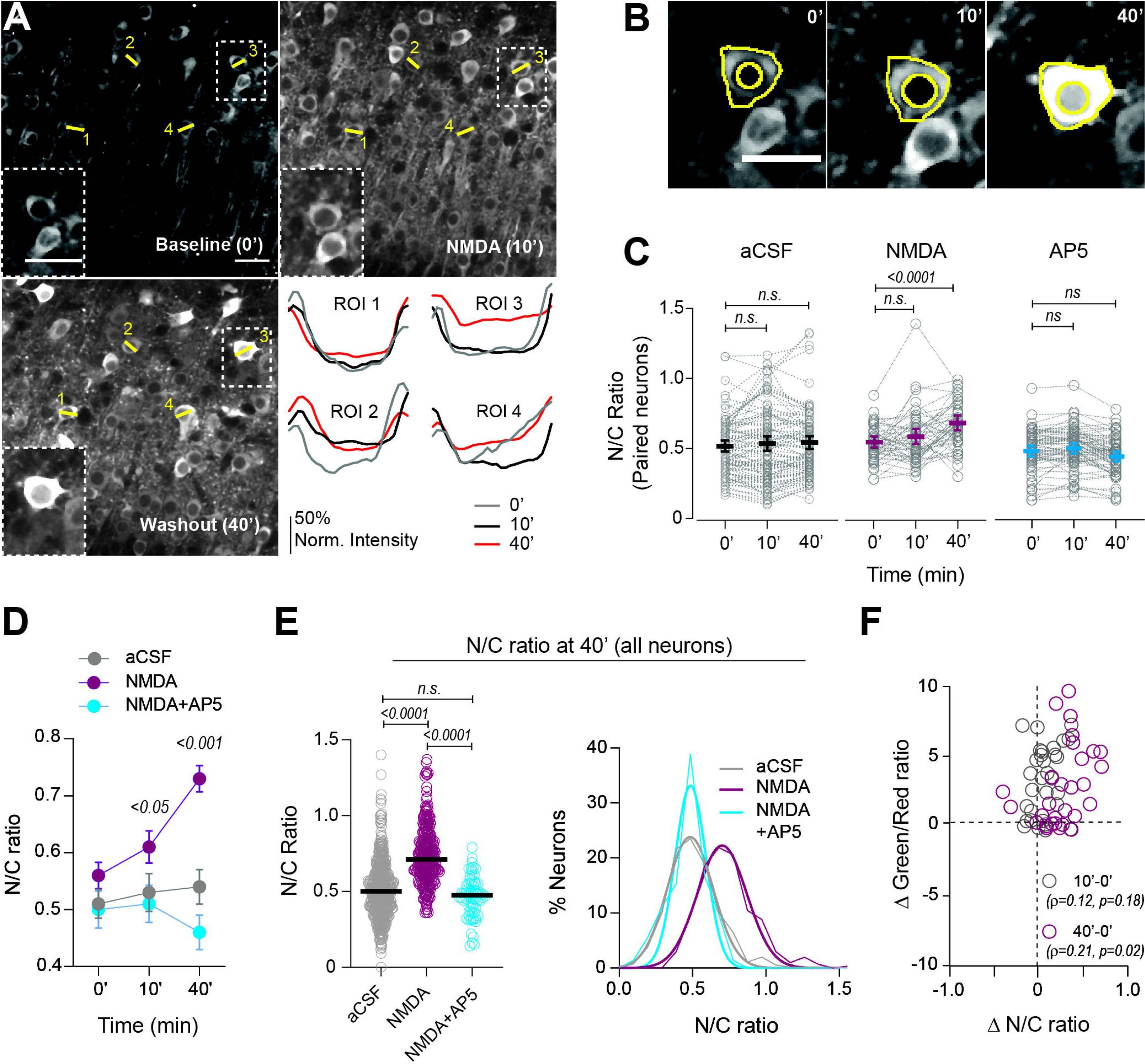
NMDA-mediated excitotoxic injury causes an increase in nuclear GCaMP6s fluorescence in neocortical neurons. **(A)** Neurons expressing GCaMP6s during baseline (0’), NMDA perfusion (10’), and washout (40’). *Inserts:* single magnified neuron showing long-lasting edema and increased nuclear and somatic GCaMP6s signal following NMDA application. Due to the swelling of extracellular space, one of the neurons in the box is out of the imaging plane. *Bottom right:* Representative traces showing normalized GCaMP6s intensity across linear ROIs drawn across somatic and nuclear compartments at multiple time points. **(B)** Overlays of nuclear ROIs (circular ROI at the centroid) and cytosolic ROIs (XOR: ANMAF-generated ROI minus nuclear ROI) used to generate compartmentalized Ca^2+^ signal intensities.**(C)** Matched comparisons of nuclear to cytosolic Ca^2+^ ratios (N/C ratios) across time in multiple conditions. RM one-way ANOVA with Dunnett’s post-test, F (1.94, 93) =12.8, p<0.0001, for aCSF alone: Freidman’s test with Dunn’s post-test, p=0.12, for AP5: Freidman’s test with Dunn’s post-test, p=0.032, n same as **Fig. 1G**. **(D)** Comparison between groups shows an increased N/C ratio at 10’ followed by a further increase at 40’. Two-way ANOVA with Tukey’s post-test, Interaction: F (4, 1044) =12.6, p<0.0001, Treatment: F (2, 1044) = 67.8, p<0.0001, Time: F (2, 1044) = 8.6, p=0.0002, n as **Fig. 1G**. **(E)** *Left:* Changes in N/C ratios using the total number of neurons at 40’ after NMDA application, rescued by AP5 application. Kruskal Wallis test with Dunn’s post-test, p<0.0001, n same as **Fig. 1I**. *Right:* distribution of N/C ratios at 40’ (all neurons). **(F)** Significant Spearman correlation between Δ Green/Red ratio and Δ N/C ratios at 40’, but not at 10’ (n=54). Data represented as mean ± 95% CI (parametric data) or median ± IQR (non-parametric data). Scale bar: 50 µm. See also **Fig. S2-S5**.

Since neurons undergo trauma during brain slice preparation (Dzhala et al., 2012), we evaluated if the N/C elevation following an excitotoxic insult also occurs *in vivo.* We performed multiphoton imaging of anesthetized Thy1-GCaMP6s pups (P11-17) at baseline, during the application of an NMDA-containing saline solution (500 µM, 10 min) over the craniotomy site and after washout (**Fig. 3A**). As seen in acute brain slices, NMDA caused significant and prolonged neuronal swelling and increased the N/C ratio *in vivo* (**Fig. 3B-D, Fig. S4F; Video S2**). Hence, nuclear translocation of GCaMP6s induced by excessive NMDAR activation, occurs *in vivo* and in acute brain slices alike.

**Fig. 3:**
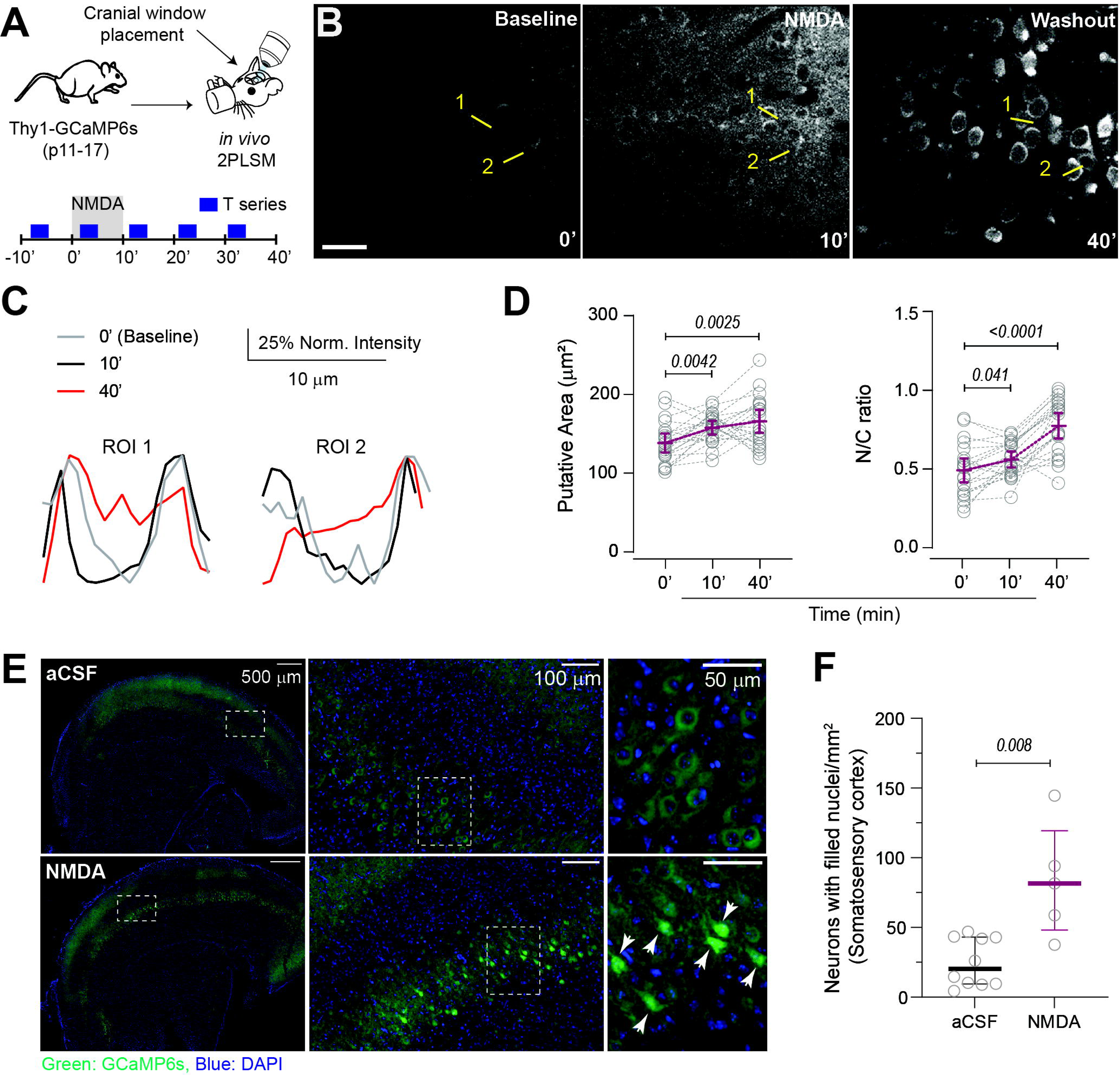
Excitotoxicity-induced nuclear translocation of GCaMP6s fluorescence *in vivo* and post-fixed brain tissue. **(A)** Schematic depicting cranial window placement and *in vivo* 2PLSM. **(B)** Neurons expressing GCaMP6s during baseline (0’), NMDA perfusion (10’), and washout (40’) *in vivo* (**Video S2**). **(C)** Normalized GCaMP6s intensity across linear ROIs drawn across somatic and nuclear compartments at multiple time points from **3B**. **(D)** *Left*: putative areas in matched neurons enlarged at 10’ and 40’ after NMDA. RM one-way ANOVA with Dunnett’s post-test, F (1.79, 35.8)=8.6, p=0.0013, n (mice: matched neurons)=4:24. *Right*: The N/C ratios (matched neurons) increased at 10’ and more so at 40’. RM one-way ANOVA with Dunnett’s post-test, F (1.86, 37.1) =46.3, p<0.0001, n=4:24. **(E)** Post-fixed slices (4% PFA) representing aCSF control (*top*) and NMDA treatment (*bottom*) showing GCaMP6s signal in green and nuclear DAPI staining in blue. Insets, higher magnification. Arrowheads: neurons with filled nuclei. **(F)** Increased number of neurons with filled nuclei after NMDA treatment. Mann-Whitney test, n (slices): aCSF=10, NMDA=6. Data represented as mean ± 95% CI (parametric data) or median ± IQR (non-parametric data). Scale bar: 50 µm **(B)** and 100 µm **(E and F)**.

Next, we performed control experiments to evaluate if the observed N/C elevation is an artifact caused by the possible saturation of the GCaMP6s signal from the intense neuronal stimulation and massive Ca^2+^ influx during NMDA application. First, we evaluated the N/C ratio change using the isosbestic point of GCaMP6s (810 nm, Ca^2+^-independent two-photon excitation) in acute brain slices (Akam and Walton, 2019). Electrical stimulus-evoked responses increased the GCaMP6s signal at 920nm excitation and diminished fluorescence at isosbestic excitation (**Fig. S5A, B; Video S3**). NMDA caused neuronal swelling in the same slices using both excitation wavelengths (**Fig. S5C, D**). More importantly, the elevation of the N/C ratio was observed in the same slices after NMDA application using both excitation wavelengths (**Fig. S5E**). Thus, we conclude that the increase in N/C is not an artifact of signal saturation but a physiological anomaly.

Second, depending on the laser intensity and duration of exposure, multiphoton imaging can induce varying degrees of phototoxicity (Peng et al., 2019), which could contribute to elevated N/C ratios. To address this, we applied NMDA (30 µM, 10 min) to acute brain slices, and instead of undergoing multiphoton imaging, they were fixed ~2 hours after NMDA application and embedded for cryo-sectioning. The neuronal GCaMP6s signal was imaged using epifluorescence microscopy (**Fig. 3E**). We again observed a marked elevation in the number of neurons with GCaMP-filled nuclei in NMDA-treated sections compared to the aCSF control (**Fig. 3F**). Together, these results conclusively indicate that the elevation in the N/C ratio is due to the entry of GCaMP6s into neuronal nuclei.

### Different insults increase the N/C ratio in neocortical neurons during early development

Robust NMDA receptor activation occurs during prolonged refractory seizures or asphyxia/anoxic depolarizations (Pietrobon and Moskowitz, 2014; Rothman and Olney, 1987; Zeiler et al., 2014), which can be cytotoxic (Fricker et al., 2018; Zhou et al., 2013) or result in neuronal death (Sadasivan et al., 2010). Thus, we evaluated if nuclear GCaMP6s translocation, observed during brief NMDA exposure, also occurs with other excitotoxic insults. First, we used repeated short NMDA puffs (3 × 100 ms) to mimic synchronous network stimulation. The NMDA puffs evoked reproducible Ca^2+^ transients in Thy1-GCaMP6s acute brain slices (P8-12 neocortex, **Fig. 4A, B; Video S4**). Neurons did not swell after multiple NMDA puffs, yet the N/C ratio increased with repeated stimulations, becoming significantly high after 8-10 stimuli (**Fig. 4C**).

**Fig. 4:**
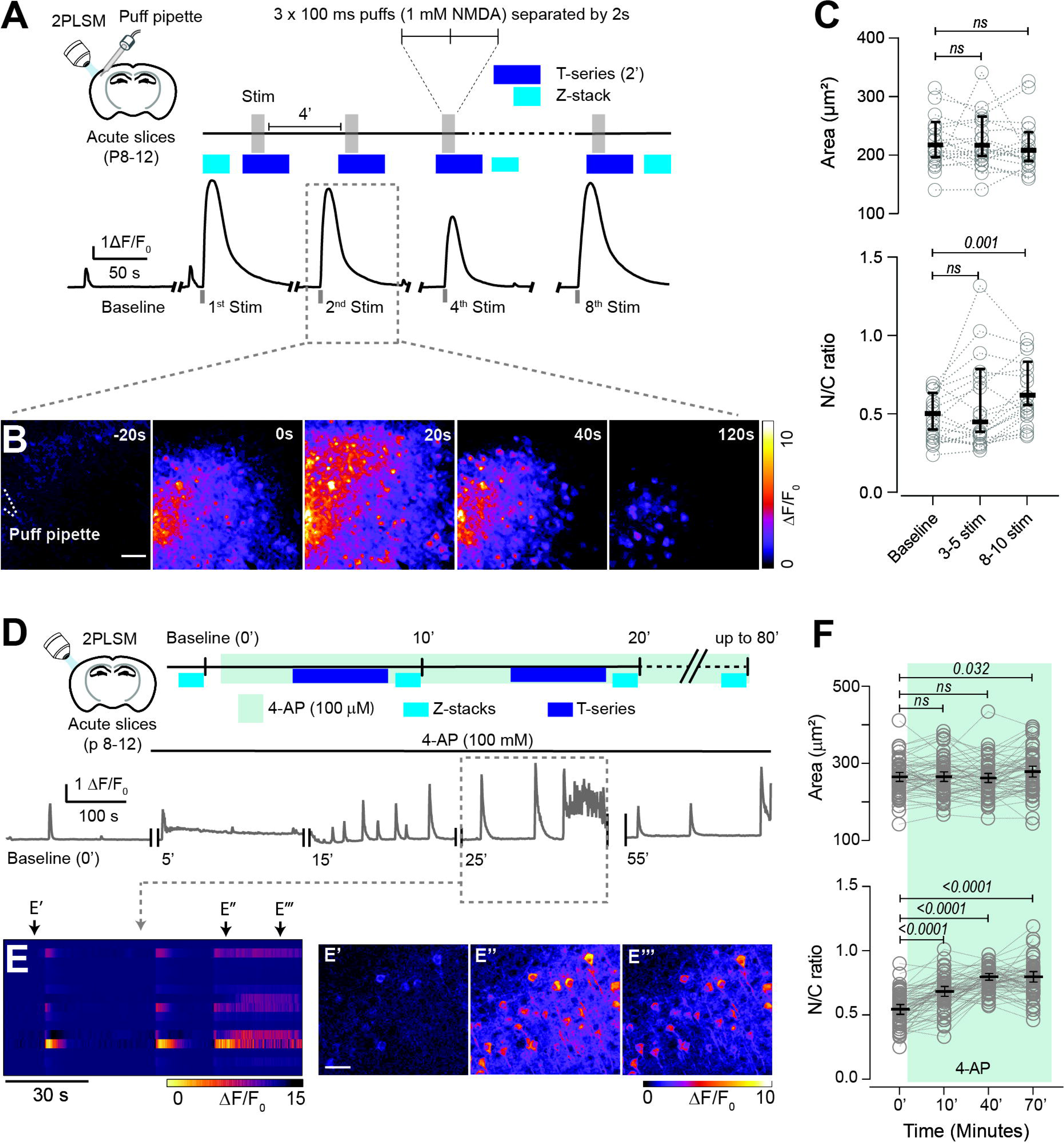
GCaMP6s nuclear translocation following repeated network stimulation and seizure-like events in the neocortical slices. **(A)** *Top*: experimental design. *Bottom:* Stimulus evoked whole-field Ca^2+^ transients. **(B)** Real-time spatial propagation of a single evoked Ca^2+^ transient (**Video S4**). **(C)** *Top*: No increase in the neuronal area following multiple stimuli. Friedman’s test with Dunn’s post-test, p=0.7, (mice: slices: neurons) n=3:4:26. *Bottom*: N/C ratio is unchanged after 3-5 stimuli but significantly increased after 8 to 10 stimuli. Friedman test with Dunn’s post-test, p=0.0013, n=3:4:26. **(D)** *Top*: experimental design. *Bottom*: Representative whole-field trace of Ca^2+^ transients (ΔF/F_0_) during seizure-like activity during 4-AP perfusion (**Video S5**). **(E)** *Left*: neuronal synchronized Ca^2+^ activity heatmap (ΔF/F_0_) during 4-AP (**E*’***, 21 neurons). *Right:* elevation of GCaMP6s signal in neurons before and during a seizure-like event. **(F)** *Top:* neurons swell after prolonged 4-AP perfusion. RM one-way ANOVA with Dunnett’s post-test, F (2.92, 175)=4.11, p=0.0081, n=3:5:64. *Bottom*: sustained increase in N/C ratio beginning 10’ after 4-AP, lasting up to 70’. RM one-way ANOVA with Dunnett’s post-test, F(2.83, 170)=80.5, p<0.0001, n=3:5:64. Data represented as mean ± 95% CI (parametric data) or median ± IQR (non-parametric data). Scale bar: 50 µm.

Second, we measured the neuronal N/C ratio during seizure-like activity in P8-12 neocortical slices perfused with the K^+^ channel blocker 4-Aminopyridine (4-AP) for 70-80 minutes while performing multiphoton imaging (**Fig. 4D**). We observed synchronous, prolonged Ca^2+^ spikes with 4-AP (**Fig. 4E, Video S5**). Z-stack images acquired during inter-ictal periods demonstrated a significant increase in the N/C ratio in neurons starting 10 min after seizure-like activity lasting throughout the experiment (~70 minutes, **Fig. 4F**). Early on, we did not observe persistent neuronal swelling. However, it was apparent at 70 minutes (**Fig. 4F**).

Third, we tested if oxygen-glucose deprivation (OGD) leads to a high neuronal N/C ratio. OGD is a hallmark feature of neurological conditions like ischemic-hemorrhagic stroke, brain trauma, and asphyxiation-hypoxia (Hartings et al., 2017; Weber and Taylor, 1994). We induced prolonged or brief OGD (reperfusion) in Thy1-GCaMP6s brain slices while performing multiphoton imaging (**Fig. S6A**). The effect of OGD was confirmed by the induction of anoxic depolarizations (AD) spreading across the neocortex (**Fig. S6B, Video S6**) (Anderson et al., 2005; Juzekaeva et al., 2020). Prolonged OGD caused neuronal swelling and a return to baseline after reperfusion, similar to our previous publication using a [Cl^-^]_i_ biosensor (Takezawa et al., 2023). Prolonged and brief OGD caused a robust and significant elevation in the neuronal N/C ratio (**Fig. S6D, E**). Although brief OGD caused no long-lasting swelling, the elevation of the N/C ratio during reperfusion was significantly larger than that caused by prolonged OGD (**Fig. S6F**), suggesting that nuclear translocation of macromolecules, like GCaMP, may represent a form of reperfusion injury (Fellman and Raivio, 1997; Ryou and Mallet, 2018). These data confirm that nuclear translocation of GCaMP is a robust and common phenotype following different forms of pathological neuronal hyperexcitability and can occur independently from neuronal swelling.

### NMDA-induced excitotoxicity increases nuclear pore size in neocortical neurons

We next evaluated the process of GCaMP6s translocation to the nucleus during excessive neuronal excitation. Bidirectional trafficking across the nuclear envelope of small (20-40 kD) and large molecules (> 40 kD) occurs through passive diffusion or by receptor-facilitated transport through the nuclear pore complex (NPC), respectively (Miao and Schulten, 2009; Wente and Rout, 2010). Nuclear transport is regulated by the endoplasmic reticulum and nuclear envelope Ca^2+^ stores and modulated by the neuronal activity (Sarma and Yang, 2011). Prior studies in cultures and transgenic mice demonstrated the degradation of the nuclear membrane and an increase in NPC size caused by prolonged (several hours to days) exposure to cytotoxic insults (Bano et al., 2010; Faleiro and Lazebnik, 2000; Sugiyama et al., 2017). We investigated whether an increase in NPC size underlies the nuclear translocation of GCaMP6s following brief excitotoxic insults reminiscent of seizures or stroke. After confirming NMDA-induced elevation of the N/C ratio using multiphoton imaging, the same Thy1-GCaMP6s slices were post-fixed, resin-embedded, and stained to prepare ultra-thin sections (~70 nm) for transmitted electron microscopy (**Fig. 5A**). The entire neuronal nuclei were imaged at low magnification (~6,000x), followed by high-magnification images of the nuclear envelope containing nuclear pores (~25,000x, **Fig. 5B**). The nuclear pore sizes were quantified as the distance between the first and last electron density peaks across a linear ROI spanning the entire pore. We found that the average nuclear pore size increased by ~29% following NMDA treatment (**Fig. 5C**), with a significant increase in the total pore size distribution (**Fig. 5D**). There was no change in the density of pores per µm, nuclear area, or nuclear perimeter (**Fig. 5E-F and I**) between aCSF and NMDA-treated neurons. Interestingly, nuclear circularity was reduced with NMDA (**Fig. 5G-H**), possibly due to an increase in activity-dependent ER invaginations into the nuclear membrane (Mozolewski et al., 2021; Oliveira et al., 2014). Therefore, these data suggest that excitotoxicity-induced increase in nuclear pore size and the resulting increase in the nuclear envelope permeability allows the translocation of a large cytoplasmatic protein, in this case, GCaMP6s, into the neuronal nucleus.

**Fig. 5:**
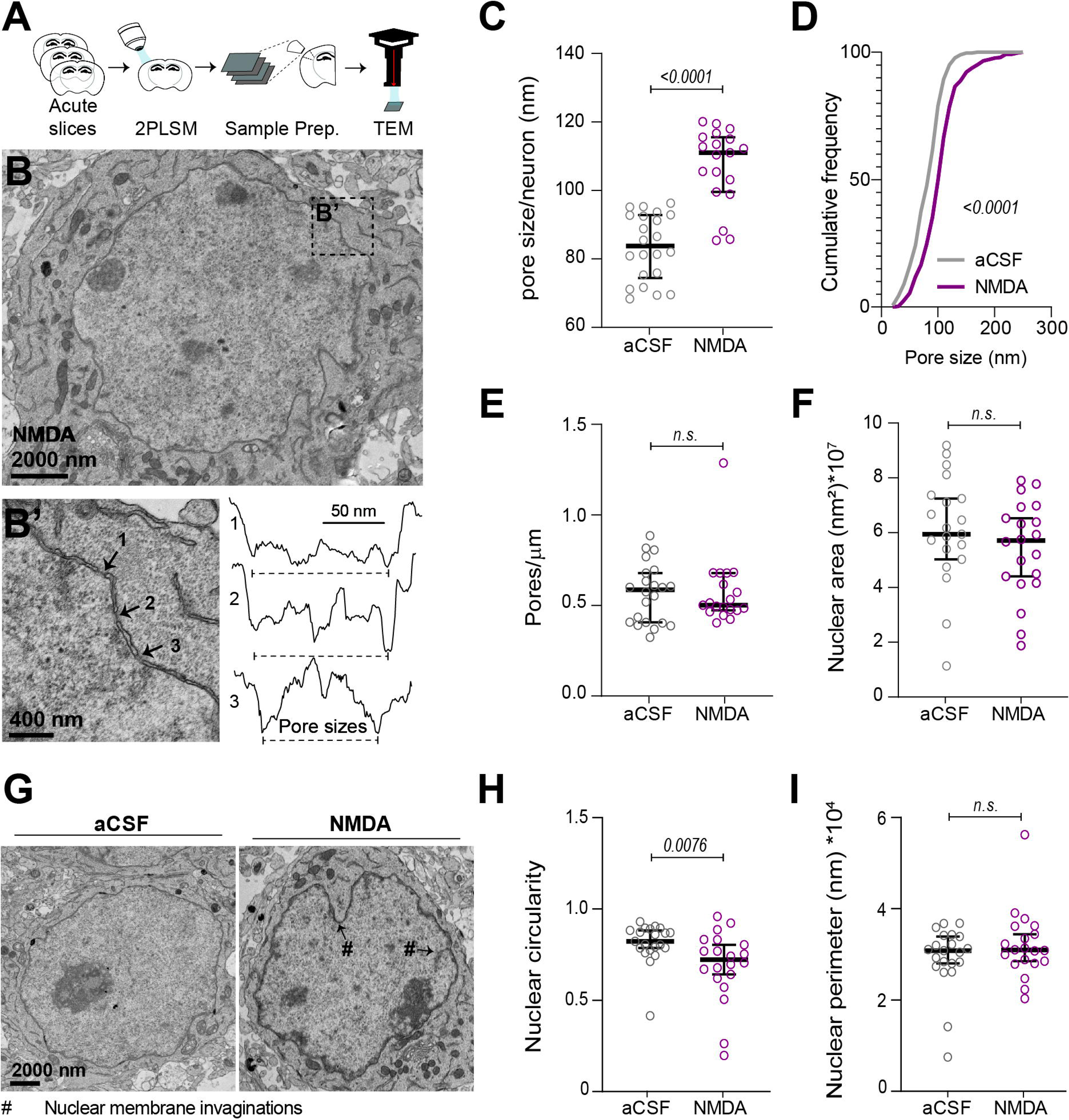
NMDA-excitotoxicity renders nuclear membranes more permeable by increasing the nuclear pore size. **(A)** Experimental design. **(B)** Transmitted electron microscopy (TEM) image of an entire neuronal nucleus (left) along with a magnification showing multiple nuclear pores (**B’**, 25,000x, arrowheads) along with normalized electron density across linear ROIs (**B’** *right*) drawn over the nuclear pores and adjacent membrane. Pore size was calculated as the distance between the first and last peaks. **(C)** NMDA significantly increased the average neuronal nuclear pore size. Mann-Whitney test, p<0.0001, n (slices: tissue samples: nuclei): aCSF=2:4:22, NMDA=2:3:19. **(D)** NMDA shifted the pore size distribution without changing **(E)** pore density or **(F)** nuclear area. Two-sample KS test, n (total pores): aCSF=343, NMDA=406. **(G)** TEM images showing nuclear invaginations (marked as #) following aCSF alone and NMDA treatments. **(H)** Nuclear circularity decreased with NMDA due to increased nuclear membrane invaginations (unpaired t-test) with no significant change in **(I)** nuclear perimeter (Mann-Whitney test). Data represented as mean ± 95% CI (parametric data) or median ± IQR (non-parametric data).

### Mitigating neuronal swelling during NMDA-mediated excitotoxicity does not prevent the increase in the N/C ratio

We next asked if the NMDA-mediated increase in neuronal N/C is preventable by mitigating neuronal swelling. Neuronal swelling is primarily mediated by the co-transport of water molecules accompanying Na^+^ and Cl^-^ influx (Andrew et al., 1997; Glykys et al., 2017; Liang et al., 2007; Rungta et al., 2015; Takezawa et al., 2023). While osmotic agents and surgical treatments mitigate brain swelling in the clinic, the prognoses remain poor for these patients (Ferrari et al., 2010; Hutchinson et al., 2016; Kolias et al., 2022). The poor clinical outcomes suggest that alternative cellular consequences of excitotoxic injury (e.g., excessive NMDA receptor-mediated Ca^2+^ influx) may contribute to the harmful effects independent of neuronal swelling. We tested if the elevated N/C ratio following brief NMDAR activation could occur independently of neuronal swelling by mitigating neuronal edema using hypertonic aCSF (+40 mOsm) (Andrew et al., 1997) as an osmotic agent.

First, we tested if osmotic perturbations alone could change neuronal N/C ratios. We treated Thy1-GCaMP6s slices with hypertonic or hypotonic aCSF that induced transient shrinkage or swelling in neocortical neurons, respectively (**Fig. S7A**). We found that hypertonic or hypotonic aCSF did not affect the N/C ratio (**Fig. S7B-C**). Next, we induced NMDA-mediated cytotoxic edema in the presence of hypertonic aCSF (+40 mOsm) to mitigate neuronal swelling. Although we observed neuronal swelling immediately after NMDA application, persistent swelling was decreased by hypertonic aCSF, and neuronal volumes returned to baseline by 40 min (**Fig. 6A, B**). The proportion of swollen neurons also decreased in hypertonic aCSF at 40 min (**Fig. 6E**). We confirmed the mitigation of NMDA-mediated neuronal swelling and dendritic beading by hypertonic aCSF by imaging YFP-expressing neurons (Thy1-H-YFP mice, **Fig. S8)**. Critically, while hypertonic aCSF decreased neuronal swelling, it did not prevent the NMDA-mediated elevation of neuronal N/C ratios (**Fig. 6C, D**). These results suggest that abnormal nuclear translocation and elevated N/C ratios of cytosolic proteins, in this instance GCaMP6s, may represent an early marker of a cytotoxic mechanism that can also occur independently from neuronal swelling.

**Fig. 6:**
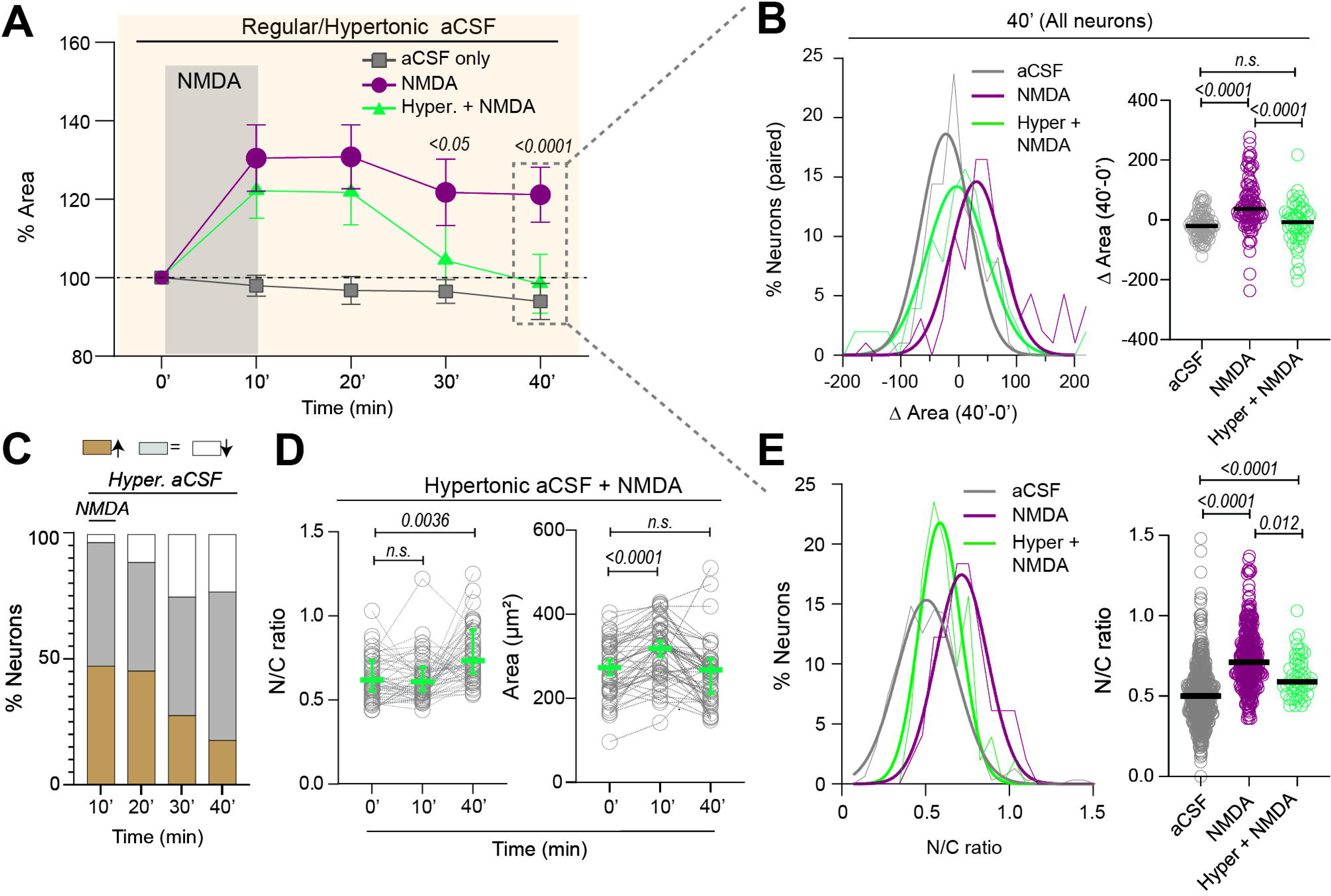
Mitigating neuronal excitotoxic edema does not prevent the persistent increase in the N/C ratio. **(A)** NMDA-induced long-term swelling is rescued with hypertonic aCSF (+40 mOsm, green) at 30’ and 40’. Two-way ANOVA with Tukey’s post-test, F(8, 915)=8.78, p<0.0001, n(mice: slices: neurons) for aCSF= 4:6:75, for NMDA= 5:8:60, for hypertonic aCSF + NMDA= 3:5:54. **(B)** *Left:* difference in areas distribution at 40’ (40’ minus 0’) in aCSF, after 10’ NMDA, and with hypertonic aCSF. *Right:* Hypertonic aCSF Δ Areas at 40’ (40’-0’). Kruskal-Wallis test with Dunn’s post-test, p<0.0001, aCSF= 312, NMDA= 234, Hypertonic aCSF + NMDA = 109. **(C)** Proportions (%) of neurons with a change in their areas (1*SD difference to baseline) at different time points with NMDA + hypertonic aCSF treatment. The percentage of swollen neurons was reduced with NMDA + hypertonic aCSF treatment (15.4±26.1) compared to NMDA alone (30.3±38.8; **Fig. S3C**). (**D)** Neuronal areas and N/C ratios measured from matched neurons across time. Hypertonic conditions prevented NMDA-induced neuronal swelling at 40’ but failed to prevent an increase in the N/C ratio. *Left,* RM one-way ANOVA with Dunnett’s post-test, F (1.62, 81) =18.6, p<0.001, n=3:5:54. *Right*, Friedman’s test with Dunn’s test, p=0.0002. **(E)** N/C ratio distributions from all neurons at 40’. Kruskal-Wallis test with Dunn’s post-test, *p*<0.0001. Data represented as mean ± 95% CI (parametric data) or median ± IQR (non-parametric data). See also **Fig. S7 and S8**.

### Nuclear translocation of GCaMP6s after NMDA-mediated excitotoxicity is calpain-mediated

Thus far, our data indicates that different excitotoxic insults to the developing neocortical neurons cause nuclear translocation of GCaMP6s and possibly other large macromolecules through an increased nuclear pore permeability independent of neuronal swelling. The nuclear translocation of GCaMP6s correlated with the elevation of intracellular Ca^2+^, suggesting a Ca^2+^-dependent mechanism. Calpains, a family of Ca^2+^-activated cysteine proteases, are linked to multiple pathologic conditions, including excitotoxicity and trauma-induced neurodegeneration and cell death (Bevers and Neumar, 2008; Cheng et al., 2018). Prior studies in neuronal cultures suggest that increased nuclear envelope permeability is associated with the activation of calpains (Sugiyama et al., 2017). Thus, we reasoned that calpains could play a critical role in mediating the observed increase in the N/C ratio. Therefore, we tested if the elevation of the N/C ratio that follows NMDA-excitotoxic injury could be prevented by inhibiting calpains in neocortical neurons. We incubated Thy1-GCaMP6s slices in a calpain inhibitor (MDL 28170) for at least 30 minutes before multiphoton imaging (**Fig. 7A, B**) (Thompson et al., 2010; Üçeyler et al., 2010). Surprisingly, MDL pre-treatment significantly reduced NMDA-induced persistent neuronal swelling measured 40 min after NMDA-treatment. Moreover, MDL, in the presence of hypertonic aCSF, further mitigated NMDA-induced neuronal swelling (**Fig. 7C**). More importantly, MDL prevented NMDA-induced elevation in the N/C ratio compared to vehicle-treated neurons. A similar suppression of N/C elevation was observed with MDL plus hypertonic aCSF (**Fig. 7D, E**). Next, to confirm these results *in vivo*, we injected Thy1-GCaMP6s pups (P11-17) with MDL (IP: 30mg/Kg) ~1 hr before craniotomy (**Fig. 7F**). We found that blocking calpains with MDL prevented NMDA-mediated neuronal swelling in young pups *in vivo* (at 40 min, **Fig. 7G**) and most importantly, MDL prevented the elevation of the N/C ratio compared to vehicle-treated mice (**Fig. 7G**), consistent with our acute brain slices results. These results indicate that Ca^2+^-regulated calpain activation contributes to increased permeability of the nuclear envelope and subsequent nuclear translocation of GCaMP6s following excitotoxic injury in the developing brain.

**Fig. 7:**
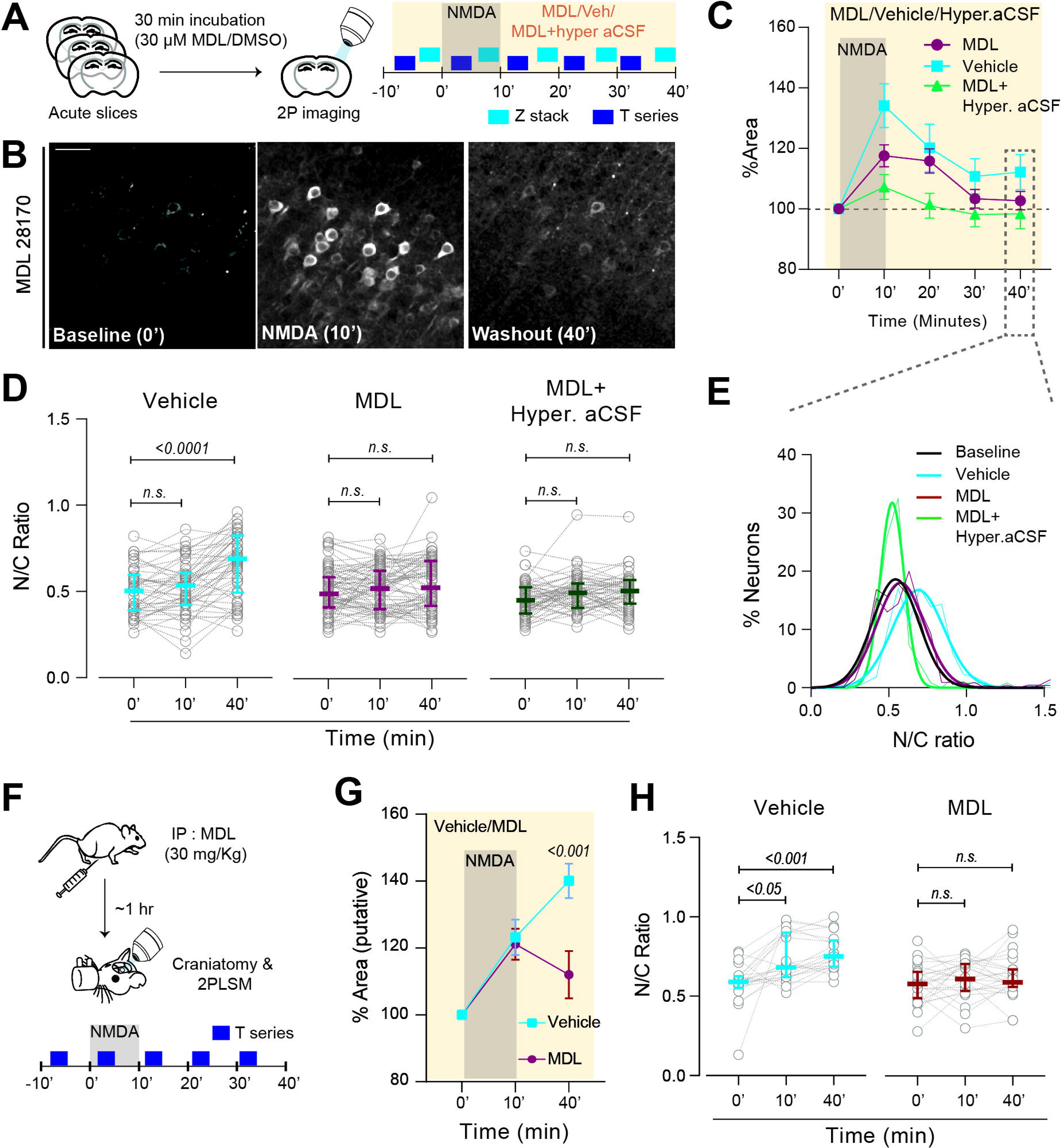
Calpain inhibition prevents nuclear translocation of GCaMP6s after NMDA-induced excitotoxic injury. **(A)** Experimental design. (**B)** Neurons expressing GCaMP6s during baseline (0’), NMDA perfusion (10’), and washout (40’) after calpain inhibition. (**C)** NMDA-induced neuronal swelling decreased with the calpain inhibitor (MDL 28170) compared to vehicle. Two-way ANOVA, Interaction: F (8, 825) =5.58, p<0.0001; Treatment: F (2, 825) =50.4, p<0.0001; Time: F (4, 825) =40, p<0.0001; Tukey’s post-test at 40’: vehicle vs MDL, p=0.004; vehicle vs MDL + hyper. aCSF, p<0.0001; n(mice: slices: neurons), vehicle=3:4:49, MDL=4:7:72. MDL + hypertonic aCSF treatment further reduced NMDA-induced swelling at 40’ (p<0.0001; n=3:4:47). (**D)** N/C ratios from matched neurons across time. Calpain inhibition prevented the increase in the N/C ratio. Friedman’s test with Dunn’s post-test, Vehicle: p<0.0001; MDL: p=0.06; MDL + Hypertonic aCSF: p=0.11. **(E)** Frequency distribution of N/C ratios from all neurons shows an increase in vehicle compared to baseline but not in MDL and MDL + hypertonic aCSF groups. **(F)** Schematic showing MDL pre-treatment (IP: 30mg/Kg) and 2P imaging *in vivo*. **(G)** NMDA-induced neuronal swelling was reduced by a single MDL dose at 40’. Two-way ANOVA with Tukey’s post-test, F(2, 132) =5.6, p=0.0046, n (mice: neurons), vehicle= 2:14, MDL= 4:18. **(H)** *In vivo* N/C ratios from matched neurons across time. MDL prevented the N/C ratio from increasing. RM one-way ANOVA with Dunnett’s post-test, vehicle: F(1.21, 24.2)=14.0, p=0.0005. MDL: F(1.7, 40.7)=0.72, p=0.47. Data represented as mean ± 95% CI (parametric data) or median ± IQR (non-parametric data). Scale bar: 50 µm.

## Discussion

Brain edema during early development contributes to lifelong neurological deficits, yet few studies evaluate the downstream consequences of excitotoxic injury at this young age. Here, we report the nuclear translocation of cytosolic GCaMP6s, a fluorescent Ca^2+^ indicator, minutes after excitotoxic insults in neocortical neurons during early brain development in acute brain slices and *in vivo*. We observed that different insults induced a simultaneous swelling and an elevation in [Ca^2+^]_i_ in neocortical neurons (**Fig. 1**), accompanied by a gradual increase in the nuclear GCaMP6s signal, which was quantified as an increase in the N/C ratio (**Fig. 2-4**). Elevated N/C ratio was associated with persistent [Ca^2+^]_i_ elevation and could occur independently of swelling (**Fig. 4, Fig. S6-7**). The nuclear entry of GCaMP6s due to excitotoxic injury was associated with larger nuclear pore sizes (**Fig. 5**). Critically, mitigating excitotoxic edema using hyperosmotic conditions alone did not normalize the elevated N/C ratio (**Fig. 6**). Yet, this elevation was prevented by inhibiting calpains (**Fig. 7**). Thus, our data demonstrates that mitigating neuronal swelling during brain injury does not prevent the abnormal translocation of cytoplasmic proteins like GCaMP6s, a marker of dysregulated nuclear function, which may underly the poor clinical outcomes associated with current brain edema treatments (Hutchinson et al., 2016; Kolias et al., 2022).

In our experiments, multiple hyperexcitability or excitotoxicity-inducing insults independently caused nuclear translocation of cytosolic GCaMP6s in a subset of neocortical neurons. GCaMPs are among the most widely used Ca^2+^ indicators due to their high sensitivity to Ca^2+^, good signal-to-noise ratio, and rapid response kinetics (Akerboom et al., 2012; Chen et al., 2012). Although their expression is restricted to the cytosol, reports suggest that GCaMPs can abnormally translocate to nuclei in certain neurons. Neurons with abnormal GCaMP nuclear accumulation often show altered Ca^2+^ dynamics and intrinsic excitability (e.g., increased firing rates) (Resendez et al., 2016; Tian et al., 2009; Yang et al., 2018). These neurons are usually discarded as “unhealthy cells” from analysis. However, the precise timing of this nuclear accumulation of GCaMPs or the underlying mechanisms are unclear. Abnormal nuclear accumulation of virally-delivered GCaMP6s and a resultant elevation in nuclear to cytosolic fluorescence ratio have been described in cultured cortical neurons a few weeks after viral infection (Yang et al., 2018). In contrast, we showed real-time nuclear translocation of GCaMP6s in neurons tracked across multiple time points minutes after exposure to brief excitotoxicity during early development. The abnormal nuclear translocation of GCaMP6s occurred following various excitotoxic insults in acute brain slices, post-fixed tissue, and in *vivo* multiphoton imaging, confirming the robustness of our findings. Our study is the first to demonstrate a calpain-mediated nuclear translocation of GCaMP6s through enlarged nuclear pores minutes after excitotoxic insults during early brain development. As nuclear pore complex degradation, which increases the leakiness of the nuclear barrier, is often observed in apoptotic or injured neurons (Bano et al., 2010; Faleiro and Lazebnik, 2000; Strasser et al., 2012), nuclear translocation of GCaMPs could be used as an early biomarker of neuronal injury.

The elevation of the N/C ratio was associated with an increase in the size of neuronal nuclear pores, suggesting altered nuclear macromolecular transport, perhaps causing dysregulation of nuclear function. Our data supports prior evidence from cultures that neuronal cytotoxic injury increases nuclear envelope permeability, resulting in the transport of cytosolically localized macromolecules into the nucleus (Ferrando-May et al., 2001; Strasser et al., 2012). The nuclear pore complexes (NPCs) mediate bidirectional transport between the cytoplasm and the nucleus. The pores form a size-selective ‘molecular sieve,’ restricting the passive diffusion of proteins larger than 40 kDa (Corbett and Silver, 1997; Miao and Schulten, 2009). Larger molecules move across the nuclear envelope through a transport receptor-facilitated translocation (Sarma and Yang, 2011; Wente and Rout, 2010). Abnormally elevated cytoplasmatic Ca^2+^ levels following prolonged activation of NMDARs can massively impair nuclear transport and disrupt multiple cellular functions (Szydlowska and Tymianski, 2010). Moreover, excessive Ca^2+^ influx following prolonged NMDAR activation can lead to calpain-mediated degradation of NPC components, increasing the nuclear membrane permeability (Norberg et al., 2008; Sugiyama et al., 2017; Yamashita et al., 2017). However, most of this experimental evidence supporting excitotoxic damage to NPCs was obtained from cultured neuronal or non-neuronal cells following prolonged exposure to excitotoxic stimuli (hours or days). Also, it is relatively unknown whether excitotoxicity alters neuronal nuclear dynamics during early developmental time points. Our study is the first to show that brief exposure to physiologically relevant excitotoxicity *in vivo* and in acute brain slices causes an increase in neuronal nuclear pore size independent of neuronal swelling. To understand the fate of the neurons with elevated N/C ratios, further work is needed to correlate the increase in the N/C ratio to known markers of cytotoxic injury and to understand how it is that not all neurons exposed to brief hyperexcitability show an elevated N/C ratio. Excitotoxicity-induced increase in nuclear membrane permeability in developing neurons can disrupt the transport of transcription factors and other regulatory proteins, resulting in an altered gene expression (Komeili and O’Shea, 2000). Additionally, Ca^2+^ is a potent regulator of gene expression, and nuclear Ca^2+^ signaling, in response to various stimuli, can facilitate or suppress gene transcription (Bading, 2013; Limbäck-Stokin et al., 2004; Papadia et al., 2005), including cell death pathways (e.g., Caspase3 or MLKL) (Zhang et al., 2007) or the suppression of transcription-dependent neuroprotective pathways (e.g., CREB/BDNF) (Hardingham et al., 1997; Hardingham and Bading, 2002). Further work is needed to identify transcriptional anomalies in developing neurons following excitotoxicity-induced persistent elevation in Ca^2+^ or increased nuclear permeability.

Inhibiting calpains prevented NMDAR-mediated nuclear translocation of GCaMP6s, both *in vivo* and in acute brain slices. Calpains are a family of calcium-dependent proteases with two major isoforms in the brain (calpain 1 and 2, also called µ- and m-calpain, respectively). Both isoforms are activated by elevated [Ca^2+^]_i_ following intense neuronal stimulation (Bevers and Neumar, 2008; Metwally et al., 2021). While both isoforms share a variety of substrates, their physiological roles can often be contradictory. Calpain-1 activation downstream of synaptic NMDARs promotes long-term hippocampal potentiation (LTP), while calpain-2 activation via extrasynaptic NMDARs can hamper synaptic potentiation and contributes to neurodegeneration (Baudry and Bi, 2016; Wang et al., 2013; Xu et al., 2009). However, their regulatory role in cell death is complementary, with both isoforms affecting a wide array of substrates that mediate necrotic, apoptotic, and autophagic cell death (Cheng et al., 2018; Müller et al., 2014; Ozaki et al., 2013; Williams et al., 2008). Moreover, multiple studies have implicated calpains in the degradation of nuclear membranes during cell death or following intense and prolonged NMDAR activation (Bano et al., 2010; Norberg et al., 2008; Sugiyama et al., 2017; Yamashita et al., 2017). Our data support these previous findings and strongly suggest that calpain-mediated degradation of the nuclear pore underlies GCaMP6s translocation into the nucleus, following physiologically relevant brief brain insults.

The current dogma recognizes cytotoxic neuronal swelling and resulting cell death as one of the leading causes of morbidity in conditions associated with severe brain edema (Liang et al., 2007; Rungta et al., 2015). Yet, using osmotic agents and surgical interventions to mitigate brain swelling continues to yield poor clinical outcomes (Hutchinson et al., 2016; Kolias et al., 2022). Swelling after intense neural activity often depends on the influx of Na^+^ predominantly through voltage-gated channels and glutamate receptors and Cl^-^ influx through various channels and transport mechanisms (Glykys et al., 2017; Rungta et al., 2015; Takezawa et al., 2023). Since neurons do not express water-permeant aquaporins, water accumulates using ionic transport through different pathways, including the cation-chloride cotransporters (Glykys et al., 2019; Takezawa et al., 2023). Cl^-^ and other ions can accumulate in neurons following intense activity, which can induce ionic overload and, in turn, alter synaptic responses and worsen neuronal intrinsic excitability (Dzhala et al., 2012, 2010; Glykys et al., 2014; Graham et al., 2022; Rahmati et al., 2021), leading to further swelling. However, neuronal swelling might not always cause cell death. Homeostatic regulatory responses can normalize the neuronal volume and ionic gradients in response to pathogenic cell swelling or shrinkage, limiting the impact that neuronal swelling may have on eventual cytotoxic injury (Hoffmann et al., 2009; Inoue et al., 2005; Lambert et al., 2008). This homeostatic neuronal volume regulation is present in developing neurons during brief OGD (Takezawa et al., 2023) and replicated in this study and during hypo- or hyperosmotic conditions (**Fig. S7**). Also, not all neurons swell or return to baseline after volume changes (**Fig. S7, 8**) (Takezawa et al., 2023). We report a novel, calpain-mediated mechanism of abnormal nuclear translocation of macromolecules independent of swelling as an early expression of neuronal injury. We speculate that this swelling-independent mechanism of neuronal injury can begin to explain why, even with adequate control of brain edema, including surgical decompression, there are no significant morbidity improvements (Hutchinson et al., 2016; Kolias et al., 2022). This clinical observation is supported by our findings that hypertonic saline improved neuronal swelling but did not decrease the N/C ratio. Significantly, blocking calpains prevented an elevation of the N/C ratio *in vivo* and in acute brain slices and decreased neuronal swelling, suggesting that blocking calpains may also aid in treating neuronal swelling.

The rodent developmental period used for our experiments (p8-12 for acute brain slices and p11-17 for *in vivo*) represents neonatal and early infancy of human development (Chini and Hanganu-Opatz, 2021; Christakis et al., 2018; Semple et al., 2013). This is a period of rapid brain development and a functional change in GABAergic neurotransmission from excitatory/shunting to inhibitory (Ben-Ari et al., 2012; Glykys et al., 2009; Tyzio et al., 2007). The excitotoxic insults used in this study (NMDA application, OGD, prolonged seizures) cause sustained neuronal depolarization and subsequent elevation in Ca^2+^ despite depolarizing or hyperpolarizing effects of GABA (Hartings et al., 2017; Juzekaeva et al., 2020; Kubista et al., 2019; Swann et al., 1993). More importantly, these excitotoxic insults at early developmental periods recapitulate perinatal brain trauma or hypoxic-ischemia (Fellman and Raivio, 1997; Ferrari et al., 2010; Mujsce et al., 1990) and helped us uncover edema-independent mechanisms of injury in neurons during early development.

In conclusion, we observed that multiple brief insults to developing neocortical neurons caused a calpain-dependent increase in the permeability of the nuclear pores, resulting in the translocation of GCaMP6s from the cytoplasm into the nucleus. This phenomenon could occur independent of neuronal swelling and can contribute to neuronal injury and subsequent poor clinical outcomes associated with posttraumatic brain edema. Additionally, nuclear translocation of fluorescent macromolecules after excitotoxic insults could be a valuable biomarker of early neuronal injury.

## Acknowledgments

We thank Jax Labs and Dr. Douglas Kim (HHMI) for the Thy1-GCaMP6s mice, and Dr. Joshua Sanes (Harvard University) for the Thy1-H-eYFP mice. pAAV-hSyn1-mRuby2-GSG-P2A-GCaMP6s-WPRE-pA was a gift from Drs. Tobias Bonhoeffer, Mark Huebener, and Tobias Rose (through Addgene). We sincerely thank Chantal Allamargot and the Central Microscopy Research Facility (CMRF) at The University of Iowa for their help with transmission electron microscopy. This work was supported by the National Institute of Health (NIH/NINDS R01NS115800) and the Iowa Neuroscience Institute. This research was partly supported by the computational resources provided by The University of Iowa, Iowa City, Iowa, and The University of Iowa Hawkeye Intellectual and Developmental Disabilities Research Center (HAWK-IDDRC) P50 HD103556.

## Author contributions

P.S. and J.G. designed the study and wrote the manuscript. P.S. and R.L. designed and performed experiments. P.S., R.L., and K.F. processed and analyzed data. J.G. provided technical direction and supervision.

## Materials and methods

### Key resource table

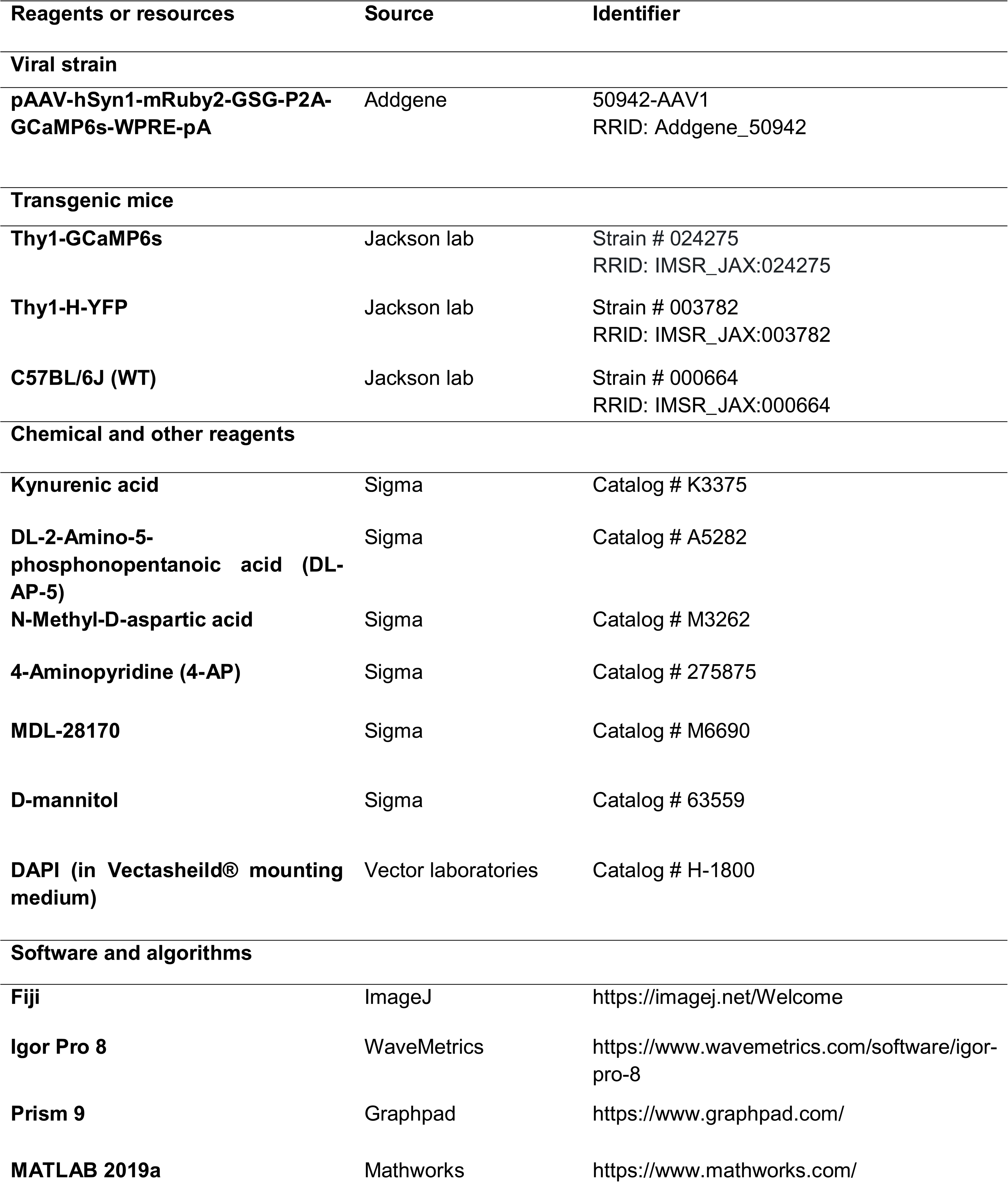

| Reagents or resources | Source | Identifier |
| --- | --- | --- |
| **Viral strain** |  |  |
| **pAAV-hSyn1-mRuby2-GSG-P2A-GCaMP6s-WPRE-pA** | Addgene | 50942-AAV1<br>RRID: Addgene_50942 |
| **Transgenic mice** |  |  |
| **Thy1-GCaMP6s** | Jackson lab | Strain # 024275<br>RRID: IMSR_JAX:024275 |
| **Thy1-H-YFP** | Jackson lab | Strain # 003782<br>RRID: IMSR_JAX:003782 |
| **C57BL/6J (WT)** | Jackson lab | Strain # 000664<br>RRID: IMSR_JAX:000664 |
| **Chemical and other reagents** |  |  |
| **Kynurenic acid** | Sigma | Catalog # K3375 |
| **DL-2-Amino-5-phosphonopentanoic acid (DL-AP-5)** | Sigma | Catalog # A5282 |
| **N-Methyl-D-aspartic acid** | Sigma | Catalog # M3262 |
| **4-Aminopyridine (4-AP)** | Sigma | Catalog # 275875 |
| **MDL-28170** | Sigma | Catalog # M6690 |
| **D-mannitol** | Sigma | Catalog # 63559 |
| **DAPI (in Vectasheild® mounting medium)** | Vector laboratories | Catalog # H-1800 |
| **Software and algorithms** |  |  |
| **Fiji** | ImageJ | https://imagej.net/Welcome |
| **Igor Pro 8** | WaveMetrics | https://www.wavemetrics.com/software/igor-pro-8 |
| **Prism 9** | Graphpad | https://www.graphpad.com/ |
| **MATLAB 2019a** | Mathworks | https://www.mathworks.com/ |

### Resource availability

#### Lead contact

Further information and requests for resources and reagents should be directed to the lead contact and corresponding author, Dr. Joseph Glykys.

#### Materials availability

This study did not use nor generate any new, unique reagents.

#### Data and code availability

The datasets and code used for image processing and data analysis in the current study have not been deposited in a public repository but are available on request.

### Experimental methods

#### Study approval and design

All experiments were conducted using a protocol approved by the Institutional Animal Care and Use Committee of The University of Iowa. Experiments were conducted on male and female pups.

#### Animals

We used transgenic mice expressing a genetically encoded calcium indicator, GCaMP6s, under the *Thy1* promoter. Acute brain slices were prepared using postnatal days 8 to 12 (P8-12) hemizygous and homozygous male and female Thy1GCaMP6s mouse pups. *In vivo* imaging experiments were performed on slightly older (P12-17) pups under anesthesia. There was no observable difference in post-NMDA neuronal swelling and nuclear translocation of GCaMP6s between male and female Thy1-GCaMP6s mice pups. Transmission electron microscopy (TEM) experiments used four slices (P11, 2 NMDA, 2 aCSF control). Immunohistochemistry (IHC) was performed on 12 brain slices from three P8-12 pups (two male, one female). All animals were housed in a temperature and humidity-controlled vivarium with free access to food and water on a 12-hour light/dark cycle.

#### Intraventricular (ICV) Viral injections

ICV viral injections were conducted on neonatal WT (C57BL/6, Jaxlab# 000664) pups (P1-2). Co-expression of GCaMP6s and mRuby2 in neocortical and hippocampal neurons was achieved by injecting pAAV-hSyn1-mRuby2-GSG-P2A-GCaMP6s one week before imaging (**Fig. 1A**; Addgene, catalog # 50942-AAV1; 1.2×10^13^ GC/mL). Animals were anesthetized using hypothermia. The anesthetized pup’s head was sterilized with a cotton swab soaked in 70% ethanol. Two µl of the virus plus 0.05% trypan blue was slowly injected into the brain (0.8-1 mm lateral from the sagittal suture, halfway between lambda and bregma) to a depth of ~3 mm using a glass micropipette. After the injection, pups were marked using a pen, allowed to recover on a heating pad, and returned to their home cage. The injected pups were visually inspected for signs of pain or distress post-surgery and were euthanized if they were found in distress. The injected pups recovered and were in good health on the day of the experiment.

#### Preparation of acute brain slices

Male and female mouse pups were anesthetized using isoflurane inhalation and decapitated per a protocol approved by the Institutional Animal Care and Use Committee of The University of Iowa. Each brain was removed and placed in ice-cold artificial cerebrospinal fluid (aCSF) containing (in mM) NaCl (116), KCl (3.3), CaCl_2_ (1.3), NaH_2_PO_4_ (1.25), NaHCO_3_ (25), and D-glucose (10) and mannitol (20) along with a high MgCl_2_ concentration (2) and kynurenic acid (2) to block glutamatergic receptors (Osmolarity: 300 mOsm). The aCSF was saturated with carbogen (95% O_2_ and 5% CO_2_) to maintain a pH of 7.3-7.4. Thick coronal sections (450 µm) containing the sensory neocortex and hippocampus were cut using a vibratome (Leica VT1000S) while submerged in aCSF. The brain slices were then placed in an interface holding chamber containing aCSF used for physiological recordings, devoid of kynurenic acid and with 1.3 mM MgCl_2_, at room temperature for 30 min. The temperature was then slowly increased and maintained at near-physiological levels (30-32°C). Slices were incubated for at least one hour before being transferred to the recording chamber.

#### Surgery for in vivo imaging

Surgical window implantation for *in vivo* imaging was performed on the day of recording at P12-17 Thy1GCaMP6s pups. Anesthesia was induced with 5% isoflurane inhalation using the SomnoSuite delivery system (Kent Scientific Corporation). Pups were mounted on a stereotactic assembly with a nose-cone mask suitable for neonatal and young mice. A circular craniotomy (~5 mm diameter) over the sensory neocortex was done using a dental drill. When removing the skull, measures were taken to reduce damage to the dura and underlying cortex (Holtmaat et al., 2009). The brain was hydrated with saline or saline-soaked surgical foam. A barrier was built around the craniotomy using cyanoacrylate glue and dental cement (Stoelting #51458 & 51456) to retain saline over the exposed brain. The pups and the stereotaxic frame were transferred to a stage for *in vivo* two-photon imaging.

#### Multiphoton imaging of brain slices

Slices were placed in a submerged chamber constantly perfused with recording aCSF maintained at 30-32°C. The location of the sensory neocortex was determined using epifluorescence. Two-photon laser scanning microscopy (2PLSM) was performed using the Bruker Ultima galvo-resonant system mounted on an Olympus BX51WIF upright microscope body with a water immersion objective (20X, 1.0 N.A.). A single Ti: sapphire tunable laser (Mai Tai HPDS; Spectra-Physics) generated two-photon excitation (920 nm) for both GCaMP6s and mRuby2. Scanning was performed with galvo-mirrors. Emitted light was bandpass filtered at 565 nm using a dichroic mirror (T510lpxrxt, Chroma), and green and red emission wavelengths were isolated using specific filters: 525/35 nm (green) and 595/25 nm (red). Two GaAsP or multi-alkali photomultiplier tubes (PMT, Hamamatsu Photonics) were used to simultaneously acquire green and red signals. Images were acquired in Layer IV/V of the sensory neocortex. Three-dimensional stacks (3D) of raster scans in the XY plane were imaged at 2 μm intervals with a 512×512-pixel resolution. Time series acquisition of single XY planar raster scans was performed at a 256×256-pixel resolution at 2.67 frames/second. All images were acquired at 2X digital zoom. Some sections were post-fixed after 2PLSM for further analysis by IHC or TEM (see below).

#### In vivo multiphoton imaging

Pups were maintained under inhaled isoflurane anesthesia (2.5-3.5%) during the imaging session. We used a Bruker-Ultima resonant-galvo system with an identical configuration to the *in vitro* system. Imaging through the craniotomy was performed using a Nikon LWD water immersion objective (16X, 0.8 N.A.). Two-photon images were acquired using galvo-scanning at a minimum depth of 200 µm from the surface. GCaMP6s was excited at a wavelength of 920 mm, and emission was filtered and acquired using a single GaAsP PMT (Hamamatsu Photonics). Time series acquisition of single XY planar raster scans was performed at a 256x256-pixel resolution at ~2.67 frames/second speed. Movement in XY dimensions was minimal and was corrected by motion correction algorithms (ImageJ). Motion artifacts in the Z plane were identified by frames with repeated cytoarchitecture. All images were acquired at 2X digital zoom.

#### Pharmacological interventions and excitotoxic perturbations

Stock solutions were prepared for all compounds in dimethyl sulfoxide (DMSO; Sigma-Aldrich # 472301) or regular aCSF, depending on their solubility. The stocks were then diluted into aCSF. NMDA (30 µM, 10 min) was applied to induce excitotoxic injury in slices. We used 4-AP (100 µM, ~70 min) to generate seizure-like activity. Hypertonic (+40 mOsm) and hypotonic (−20 mOsm) conditions were created by adding or removing 20 mM mannitol to the regular aCSF. Whenever appropriate, vehicles contained identical DMSO concentrations to respective stock solutions of the drugs (all ≤ 1%). All drugs were perfused at 2 mL/min and took <30 seconds to reach the recording chamber. For *in vivo* experiments, MDL stock (3.8 mg/mL) was prepared in a vehicle containing 50% saline and 50% DMSO. The amount of stock (or vehicle) corresponding to the dose (30 mg/kg) was intraperitoneally injected in mice for at least 1 hour before the experiment. The total volume injected was always less than 80 µl.

#### Repeated micro-stimulation using focal NMDA puffs

Pulled glass micropipettes (0.5–1 MΩ resistance, tip size 10–20 μm) were filled with 100 mM NMDA and placed on the neocortical surface of acute brain slices. NMDA was applied as brief puffs (3psi, 100 ms) using a pressure ejector (MPPI-2, Applied Scientific Instruments). Each stimulus consisted of three brief NMDA puffs, separated by 2 seconds. Every stimulus resulted in a Ca^2+^ transient.

#### Oxygen-glucose deprivation (OGD)

To induce OGD in acute neocortical slices, D-glucose in the aCSF was replaced with D-mannitol, and the solution was saturated with 95% N_2_ (to replace O_2_) and 5% CO_2_. Before applying the OGD solution, Z-stacks and time-series images were acquired at baseline in regular oxygenated aCSF. OGD solution was applied for either 20 minutes (prolonged OGD) or 10 mins, followed by regular, oxygenated aCSF for 10 more mins (OGD + reperfusion). Z-stacks and time-series images were acquired throughout the experiment. Successful OGD induction in slices was confirmed by the appearance of slowly propagating Ca^2+^ transients associated with anoxic depolarizations or ADs.

#### Immunohistochemistry (IHC)

After *ex vivo* imaging or incubation, slices were fixed with 4% PFA in PBS at 4°C overnight. The fixed slices were then washed with PBS, dehydrated in 30% sucrose (4°C overnight), and either embedded in blocks of 15% porcine gelatin (type A) or cryopreserved in a solution containing 40% PBS, 35% ethylene glycol, and 25% glycerol. The brain blocks were cut coronally (10 μm) using a cryostat (CM3050S, Leica). Sections were permeabilized with 0.2% Triton X-100 in PBS for 30 min at room temperature. Sections were mounted on slides using a media containing 4,6-diamidino-2-phenylindole (DAPI) to visualize cellular nuclei. Slides were imaged under a 10x objective on an epifluorescence microscope (Leica SPE confocal microscope) using the following filters: blue: ex360/40, em470/40, dc400; green: ex480/40, em527/30, dc505. We used a blue filter for DAPI and green for GCaMP6s. The entire section was imaged using the Leica Application Suite X program. All images from an experimental set were recorded at 12 bits (2.20 pixels/μm) using identical fluorophore exposure times and gain for each slide. Image J was used to analyze the images. The analysis was conducted in the somatosensory region of the neocortex (average area = 1.10 mm^2^). ROIs were drawn manually for GCaMP6s.

#### Transmission Electron Microscopy (TEM)

After 2PLSM, live coronal sections were fixed in a solution containing 2% paraformaldehyde and 2% glutaraldehyde in 0.1 M Sodium Cacodylate buffer, pH 7.4. The sections were then post-fixed with 1% osmium tetroxide aqueous solution containing 1.6 % ferrocyanide and contrasted with 2.5% uranyl acetate aqueous solution. After dehydration in a graded ethanol series up to 100% and immersion in propylene oxide, sections were infiltrated with Eponate 12 resin (Ted Pella Inc, Cat#18010) and then flat-embedded between Aclar films (Ted Pella Inc, Cat# 10501) with Eponate resin, and polymerized overnight at +60°C. Chips with L4 and 5 of the sensory neocortex were selected by light-microscopic inspection, glued to blank epoxy blocks, and sectioned with an ultramicrotome (Leica microsystems). Ultrathin sections (~70 nm) were collected, mounted on 200 mesh copper grids, and stained with 5% uranyl acetate and Reynold’s lead citrate solutions. Contrasted grids were examined with an HT7800 Transmission electron microscope. Before the acquisition, L5 neurons were identified in real-time using a low-resolution CCD camera (Hitachi), and images were acquired with an AMT Nanosprint 15 digital, high-sensitivity camera system with a 5056 x 2960-pixel sensor. Low-magnification images of the whole nuclei (5 or 6,000x) and high-magnification images of the nuclear membrane segments (25,000x or more) were captured.

#### Extracellular electrophysiology in acute brain slices

Acute brain slices were placed in an interface chamber (32–34°C) and perfused with aCSF saturated with 95% O_2_ and 5% CO_2_. Glass electrodes filled with aCSF were placed in the neocortex (layers IV and V of somatosensory regions) identified using a stereomicroscope (AmScope). Extracellular field potentials were recorded using a low-noise differential amplifier (DP-311, Warner Instruments, 100x gain) and digitized at 2 kHz (IX/408, iWorx Systems Incorporated). NMDA was briefly applied to the slices (30 µM, 10 min) to induce an acute excitotoxic injury. Field potentials at baseline, during NMDA treatment, and washout were analyzed using a custom-written macro in IgorPro v8.04 (WaveMetrics).

### Measurements and statistical analysis

#### Measuring neuronal areas and intracellular Ca^2+^ levels

Z-stacks acquired using 2PLSM were background-subtracted, smoothened (median filter, radius=2), converted to maximum intensity projections (MIPs, every 10 images), and contrast-enhanced (CLAHE, ImageJ). Thus, the maximal areas for each neuron over a 20 µm depth were represented in a MIP (Glykys et al., 2019). Neuronal cell bodies in the processed MIPs were detected using an automated neuronal morphology analysis framework (ANMAF) based on a convolutional neuronal network and saved as regions of interest (ROIs) for further analysis (Tong et al., 2021). For images acquired in dual color, neuronal ROI sets were generated separately for each channel (GCaMP6s and mRuby2) using ANMAF. A custom-built macro automatically updated neuronal ROIs to the z-planes corresponding to the maximum area. Change in intracellular [Ca^2+^]_i_ was measured as the ratio of median GCaMP6s and mRuby2 intensities (Green/Red ratio) for each neuronal ROI.

#### Measuring the nuclear to cytosolic fluorescence ratio (N/C ratio)

Traditional GECIs like GCaMP6s have cytosolic localization, with most of the signal emerging from the cytosol and nuclei appearing darker in comparison. Using neuronal ROIs generated by ANMAF, we confirmed the presence of nuclei in their respective z-planes. Then, using a custom-built, automated macro, circular ROIs (15 µm in diameter) were drawn at the center of mass of all the neuronal ROIs (hereafter called nuclear ROIs), and their XY locations were adjusted (if needed). The nuclear ROIs were then subtracted from the ANMAF-generated neuronal ROIs (using the XOR function) to generate the cytosolic ROI. The ratio of mean intensities of nuclear and cytosolic ROIs was calculated for every neuron at the corresponding z-plane.

#### Measurement of neuronal Ca^2+^ transients

Image processing, demarcation of regions of interest (ROI), and collection of fluorescence traces were performed in Fiji. Fluorescence traces were analyzed in IgorPro8 (Wavemetrics) and Prism 9 (GraphPad). Raw images in time series were background-subtracted, smoothened (median filter, radius=2), and compressed into maximum intensity projections (MIPs). Circular ROIs representing neuronal somas and concentric, doughnut-shaped ROIs representing the neuropil were generated on the MIPs using a semi-automated, custom-built macro. The neuropil fluorescence was subtracted from the corresponding somatic fluorescence for each frame. To determine the characteristics of evoked Ca^2+^ responses, we performed baseline subtraction and normalization (ΔF/F_0_) of the neuropil-subtracted Ca^2+^ signal for each neuron using the following equation: (F_t_-F_0_)/F_0,_ where F_t_ is the fluorescence intensity of a given frame, and F_0_ is the average fluorescence of the first 8-10s of the trial (before micro-stimulation using NMDA puffs). To determine the characteristics of spontaneous Ca^2+^ transients, the neuropil-subtracted traces of neuronal fluorescence were transformed into Z-scores using the following equation: Z = (F_t_-F_M_)/σ, where F_t_ is the fluorescence intensity of a given frame, F_M_ is the average fluorescence of the entire trace, and σ is the standard deviation of the fluorescence for the whole trace.

#### Statistics

Experimenters were not blinded during data analysis, as the neuronal calcium activity elicited by excitotoxic insults was quite apparent. The data were acquired multiple times (0’ or baseline, 10’ or after NMDA, 20’, 30’, 40’ or washout). A subset of ANMAF-generated neuronal ROIs was matched in multiple time points using a custom-built macro (Igor 8). The normality of distributions was determined using the D’Agostino–Pearson K2 and Anderson-Darling tests. Data are presented as mean ± confidence interval (95% CI) or median ± interquartile range (IQR) based on normality distribution. As a functional correlate of intracellular Ca^2+^, ratios of median intensities of green and red channels were calculated for every neuronal ROI (Green/Red ratios) and represented as mean ± 95% CI. Matched data representing multiple time points with the same perturbation were analyzed using repeated measures one-way ANOVA with Dunnett’s test for multiple comparisons (Friedman’s test with Dunn’s multiple comparison tests for non-parametric data). Unpaired parametric data across multiple time points were analyzed using one-way ANOVA (or Kruskal-Wallis test for non-parametric data). Data with multiple time points were compared across groups using two-way ANOVA with Tukey’s test for multiple comparisons. Data points representing individual slices/cells were compared across groups using a two-tailed, unpaired t-test (parametric data) or Mann–Whitney U test (non-parametric data). Spearman rank correlation (ρ) and Pearson correlation (expressed as *r*^2^) were used for parametric and non-parametric data, respectively. Paired comparisons were performed using a two-tailed, paired t-test. Normalized (%) distributions of individual events were compared across genotypes using a two-sample Kolmogorov–Smirnov (KS) test. Statistical significance was considered at p<0.05.

