## Supplementary material for "Brief and diverse excitotoxic insults cause an increase in neuronal nuclear membrane permeability in the neonatal brain": Fig. S

**Supplementary information**

Brief excitotoxic insults cause a calpain-mediated increase in neuronal nuclear membrane permeability in the immature brain.

P. Suryavanshi, R. Langton, K. Fairhead, and J. Glykys

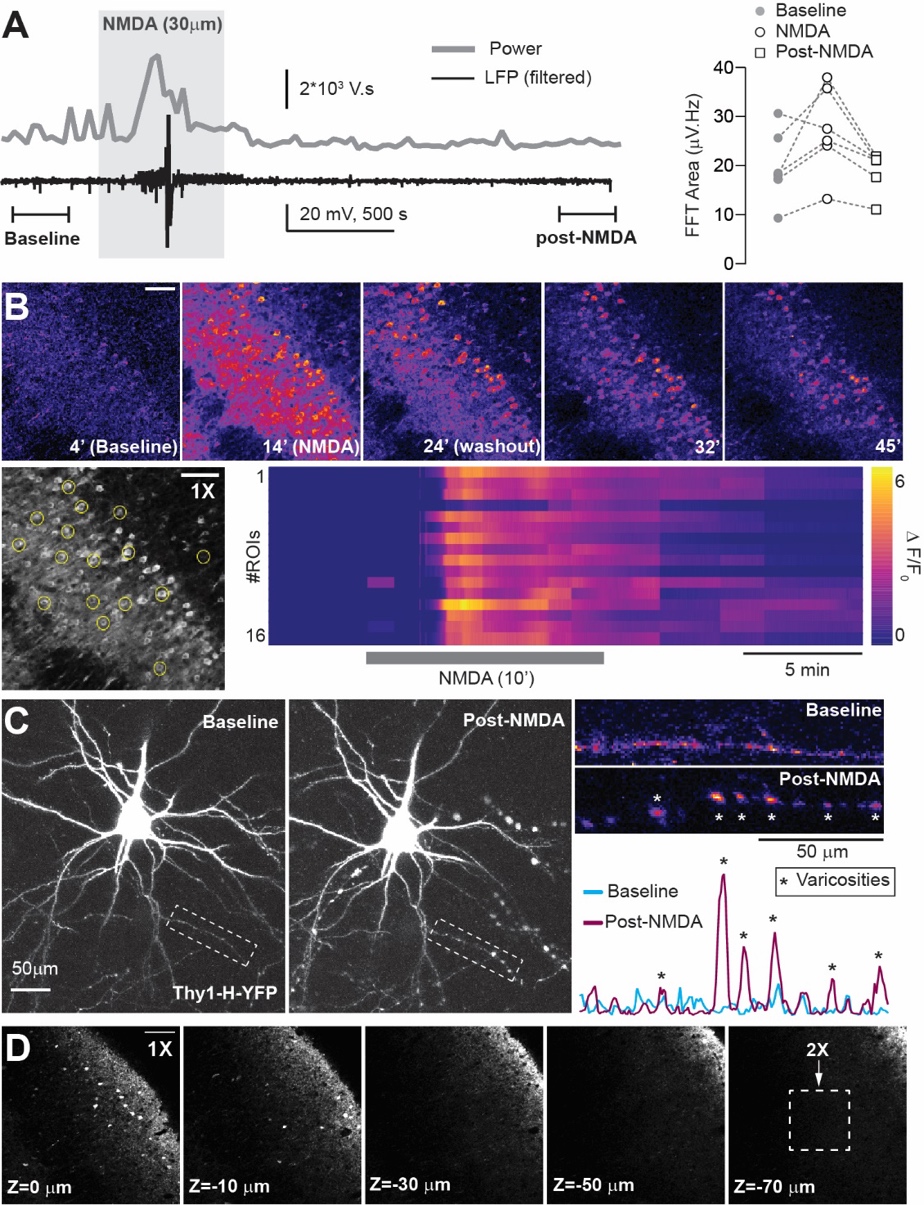

**Fig. S1. NMDA-induced hyperexcitability and neuronal injury in slices.**

**(A)** *Left:* Representative trace showing filtered local field potentials (LFP) and its FFT power calculated for a window of 30 seconds, at baseline, during, and after NMDA application (P8-12). LFP power increases during NMDA application. *Right*: Quantification of FFT areas recorded from individual neonatal slices *(n=6)* at baseline, during NMDA application, and post-NMDA. **(B)** *Top:* Representative images (Thy1-GCaMP6s) depicting NMDA-induced Ca^2+^ activity in neonatal neurons imaged for 40 mins. *Bottom:* ROI placement (left) and heatmap showing Ca^2+^ traces (right, ΔF/F_0_) generated from the ROIs. **(C)** *Left:* representative images showing NMDA-induced long-lasting dendritic injury in the form of varicosities or beading at 40’ post-NMDA (Thy1-YFP; p10-12). *Right:* Quantification of dendritic beading using kymographs (*top*) and signal intensity distribution (*bottom*). **(D)** Representative images (1X) of a Thy1-GCaMP6s slice at different depths showing the location of “injured” neurons with nuclear accumulation of GCaMP on the superficial tissue. Dashed box: normal depth of imaging area. Scale: 50 µm.

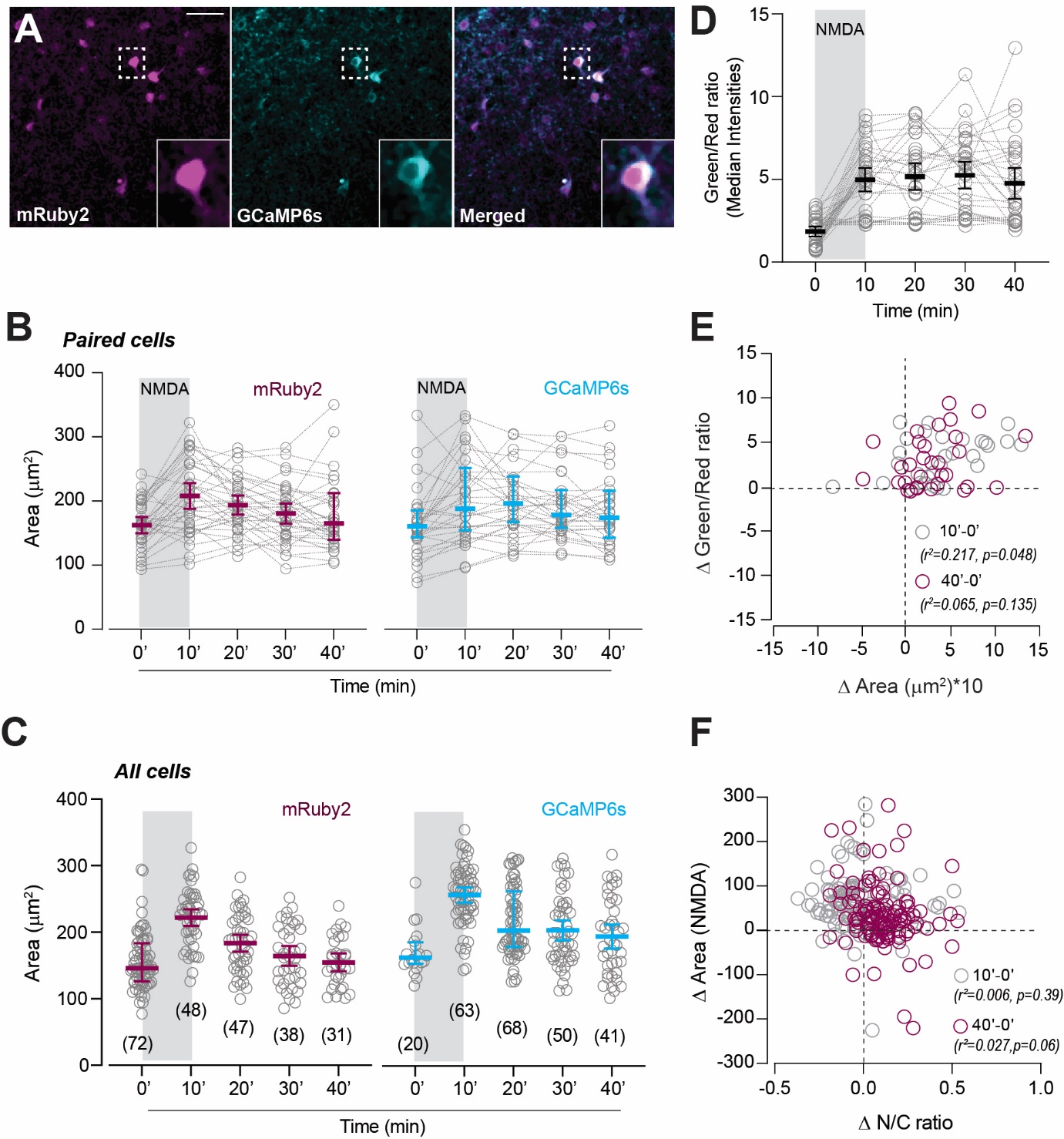

**Fig. S2. Long-term dual-color imaging of neuronal swelling.**

**(A)** mRuby2 (magenta)- and GCaMP6s (cyan)-expressing cortical neurons (P9) with a merged image. *Inset*: A single neuron at higher magnification. **(B)** NMDA-induced increase in neuronal area in matched cells across time (n=35, mRuby2; n=31, GCaMP6s). **(C)** Increase in the neuronal area in the total number of neurons. The number of ANMAF-identified neurons is shown in parentheses. **(D)** The Green/Red ratio increases across time following NMDA treatment (paired neurons, n (mice: slices: neurons)=2:3:54). **(E)** No significant Pearson correlation between the change in (Δ) Area and Δ Green/Red ratio in paired neurons at 40’ after NMDA application. **(F)** No significant correlation between Δ Area and Δ N/C ratios in paired neurons with NMDA application in slices prepared from Thy1-GCaMP6s mice (Pearson correlation, *n=87*). Data represented as mean ± 95% CI (parametric data) or median ± IQR (non-parametric data). Scale: 50 µm.

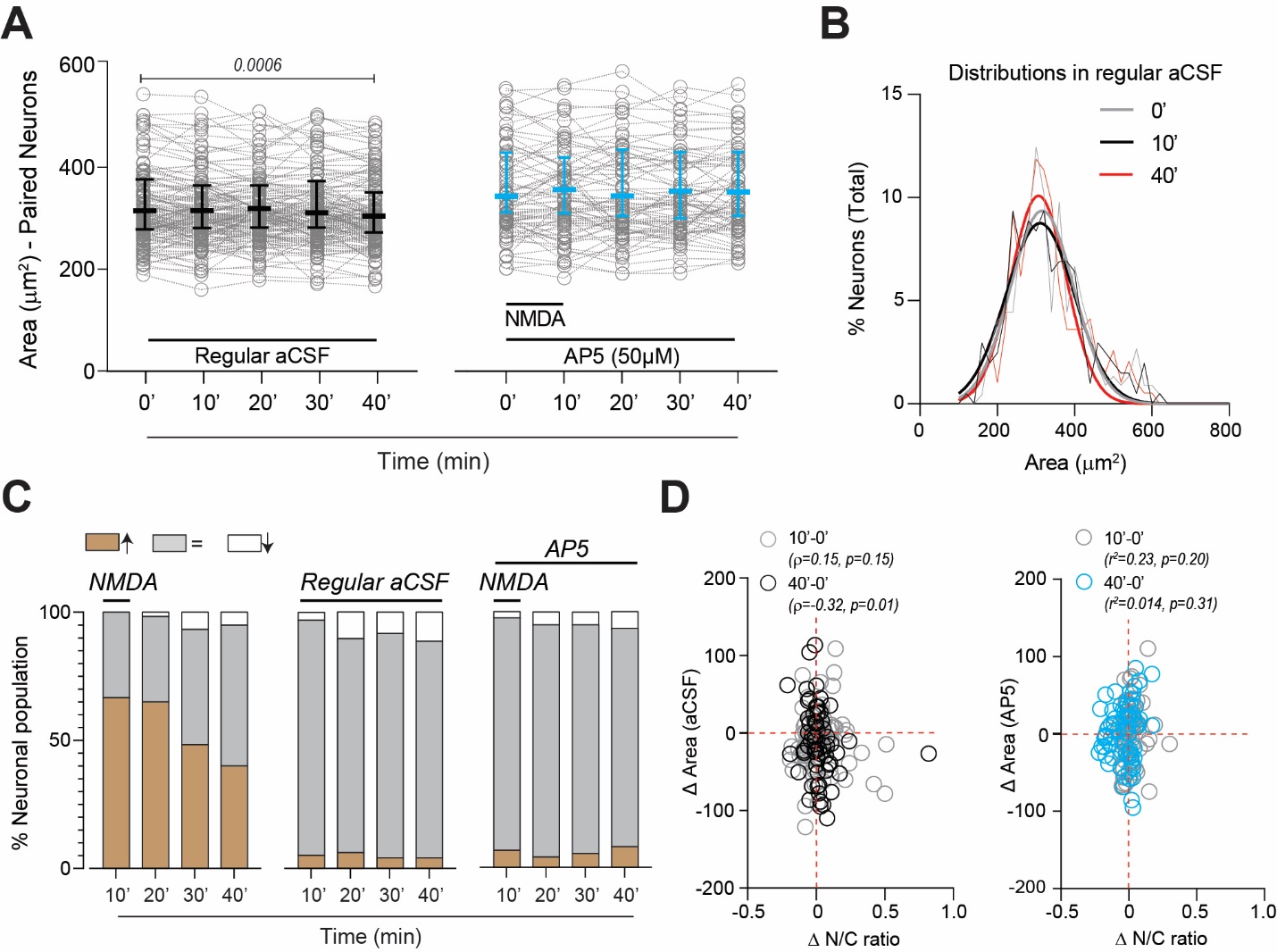

**Fig. S3. Neurons do not swell due to prolonged aCSF application or when NMDA receptors are blocked.**

**(A)** There was no significant increase in matched neuron areas across time with aCSF alone (black) or NMDA+AP5 (cyan) treatments. aCSF alone caused a small but significant shrinkage in matched neurons at 40’. RM one-way ANOVA with Dunnett’s post-test, aCSF: F(3.57, 539)=4.22, p=0.0034; NMDA+AP5: F(1.99, 143)=0.006, p=0.99. **(B)** Area distribution measured from all neurons showed no change across time (0’, 10’, and 40’) with regular aCSF. **(C)** Percentage of neurons with a change in their areas (1*SD difference from baseline) at different time points, with NMDA, aCSF alone, and NMDA+AP5. **(D)** Δ Areas were correlated with N/C ratios at 40’ with aCSF alone (*left*, Spearman correlation). No correlation existed between Δ Areas and N/C ratios at 10’ or 40’ with NMDA+AP5 (*right*, Pearson correlation). Data represented as mean ± 95% CI (parametric data) or median ± IQR (non-parametric data).

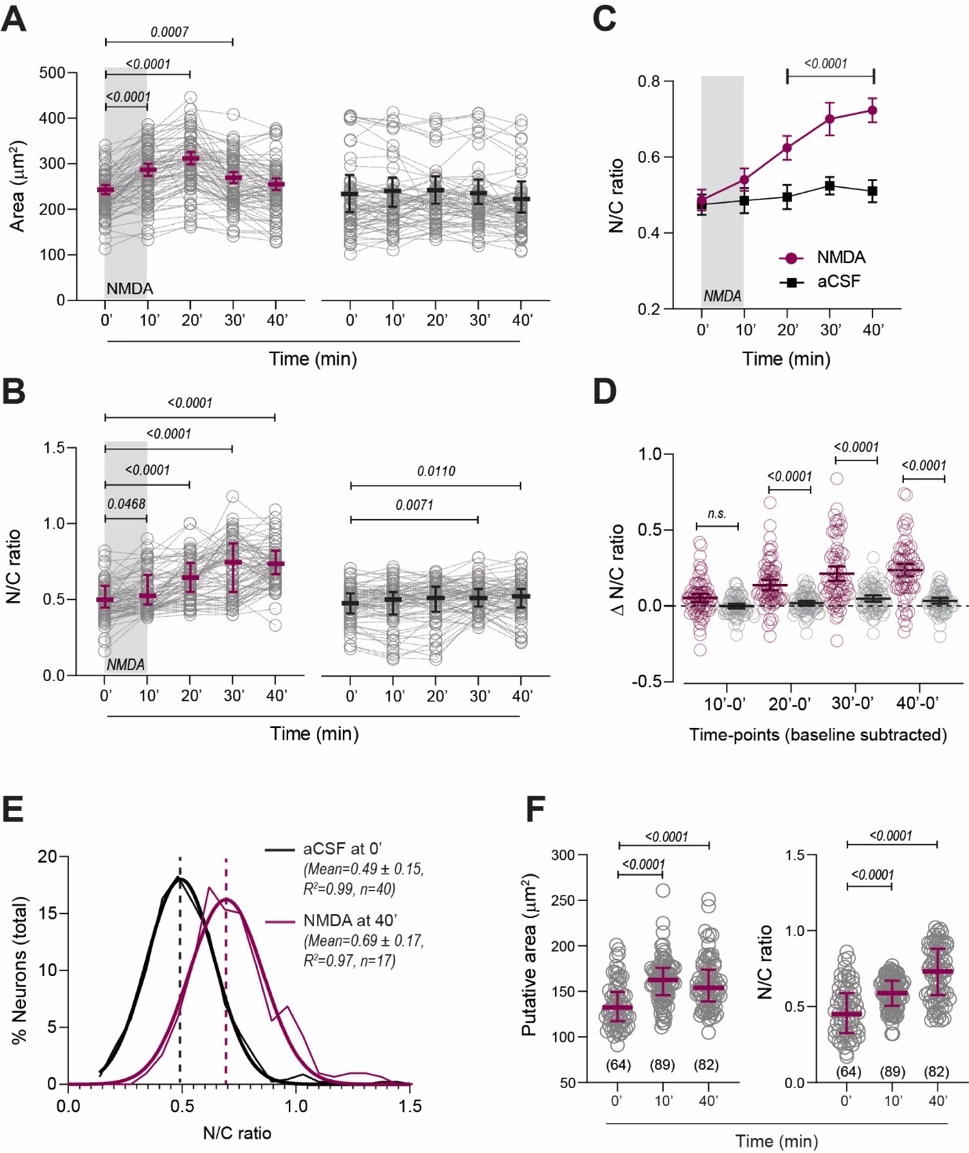

**Fig. S4.** **The neuronal N/C ratio increases shortly after NMDA treatment and remains elevated.**

**(A)** Area changes in matched neurons following NMDA (magenta) or aCSF alone (black) treatments over time. RM one-way ANOVA with Dunnett’s post-test; NMDA: F(3.05, 226)=51.8, p<0.0001; aCSF: F(3.36, 232)=3.42, p=0.014; n (mice: slices: neurons), NMDA=3:6:76, aCSF=2:4:71. For this dataset, all neuronal ROIs were drawn manually. **(B)** Changes in the N/C ratios of matched neurons following NMDA (magenta) or aCSF alone (black) treatments at different time points. Friedman test with Dunn’s post-test, NMDA: p<0.0001, aCSF: p=0.001. **(C)** N/C ratio between NMDA and aCSF treatments across time. N/C ratios significantly increase after NMDA treatment, at 20’ onwards. Two-way ANOVA, Interaction: F(4, 710) =14.1, p<0.0001; Time: F(4, 710) =29.3, p<0.0001; Treatment: F(1, 710) =139.5, p<0.0001; Sidak’s post-test for timepoints from 20’ to 40’, p<0.0001). **(D)** Change in N/C ratio relative to baseline (Δ N/C ratio) between NMDA and aCSF treatments across time. Two-way ANOVA with Sidak’s post-test, Interaction: F(3, 568) =8.7, p<0.0001; Time: F(3, 568) =22.5, p<0.0001; Treatment: F(1, 568) =152.5, p<0.0001. **(E)** N/C ratios distribution from all neurons at baseline (aCSF alone at 0’, n=40 slices) and 40’ after NMDA application (n=17 slices). **(F)** Significant increase in putative areas and N/C ratio calculated from all neurons *in vivo* following NMDA treatment*.* Kruskal-Wallis test with Dunn’s post-test, p<0.0001. Data represented as mean ± 95% CI (parametric data) or median ± IQR (non-parametric data).

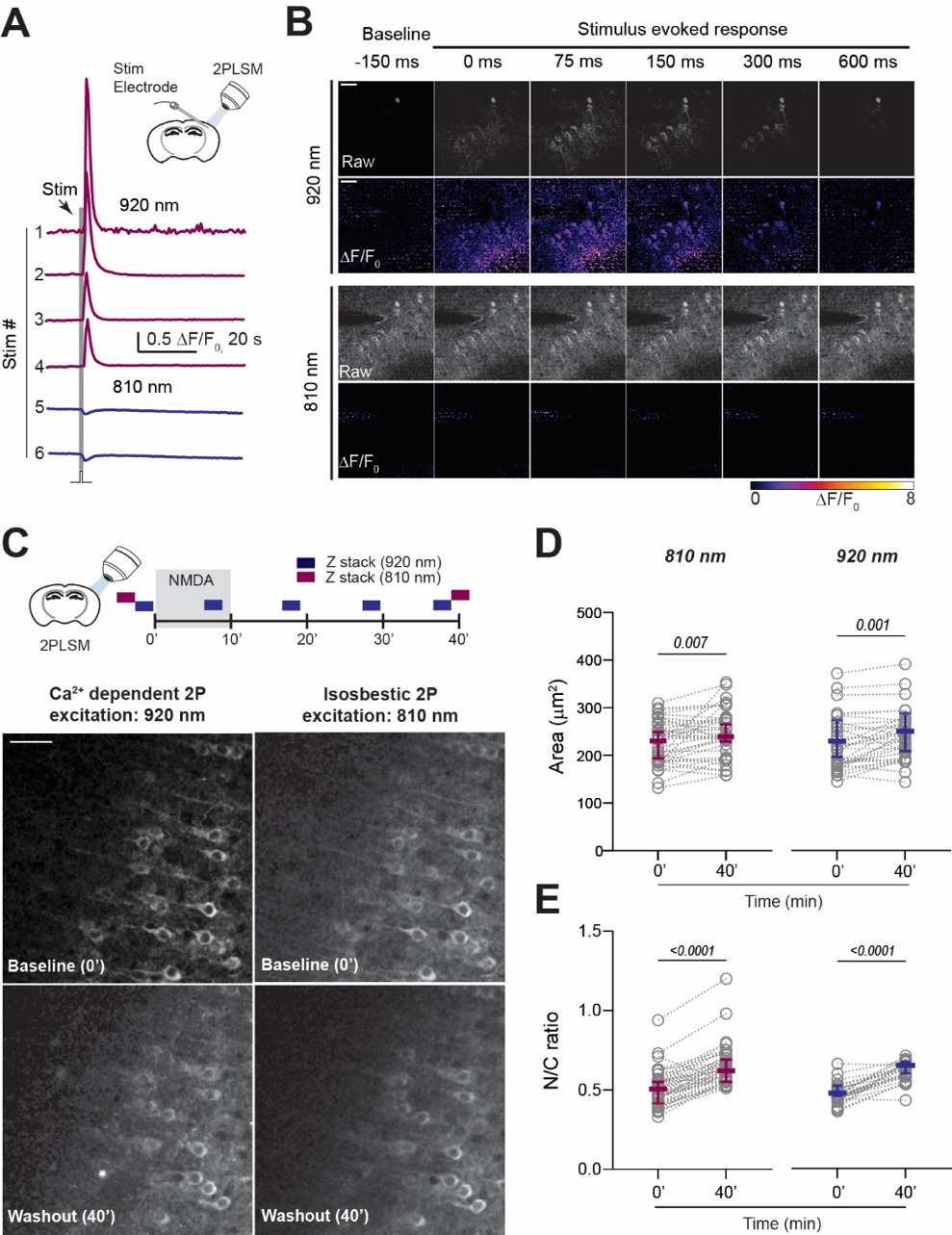

**Fig. S5. NMDA-induced nuclear translocation of GCaMP6s imaged using** **Ca^2+^ independent isosbestic excitation.**

**(A)** Representative electrical stimulus-evoked whole-field Ca^2+^ transients acquired at 920 nm and with isosbestic excitation at 810 nm. **(B)** Representative images (raw and ΔF/F_0_) depicting the progression of stimulus-evoked Ca^2+^ transients imaged at 920 nm (*top*) and 810 nm (*bottom,* **Video 3**). **(C)** *Top:* experimental design to capture NMDA-induced nuclear translocation of GCaMP6s signal at 920 nm and 810 nm excitation. *Bottom:* Representative images of GCaMP6s expressing neurons at baseline (0’) and washout (40’) after NMDA application. **(D)** Neuronal area increased 40’ after NMDA perfusion. GCaMP6s excitation: 810 nm and 920 nm. Paired t-test. **(E)** N/C ratio increased after NMDA treatment (40’), using 810 nm excitation *(left)* and 920 nm excitation *(right)*. Paired t-test; n (mice: slices: neurons) = 2:3:38 (810 nm) and n=2:3:40 (920 nm). Data represented as mean ± 95% CI (parametric data) or median ± IQR (non-parametric data). Scale bar: 50 µm.

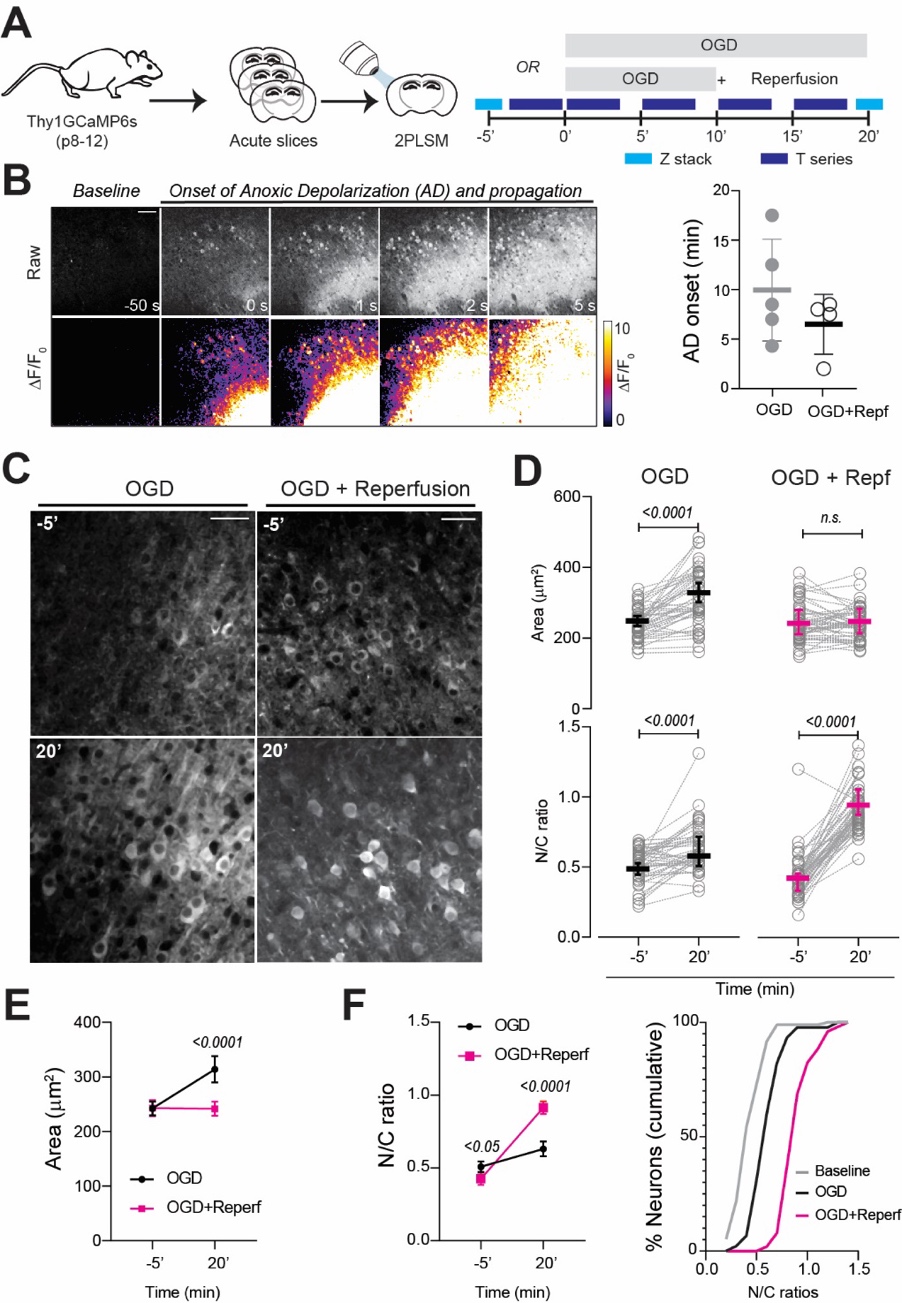

**Fig. S6. Oxygen-glucose deprivation (OGD) increases the neuronal N/C ratio in the neonatal neocortex.**

**(A)** Experimental design to induce prolonged (OGD for 20’) and brief OGD with reperfusion (OGD+reperf, 10’ each). **(B)** *Left:* representative images (raw and ΔF/F_0_) depicting real-time propagation of an anoxic depolarization (AD) event during OGD induction (**Video 6)**. *Right:* AD onset in different slices. **(C)** GCaMP6s-expressing neurons at baseline (-5’) and after induction of prolonged or brief OGD (20’). **(D)** *Top:* increase in neuronal area with prolonged OGD but not after brief OGD followed by reperfusion. *Left*, paired t-test, n =3:5:43; right, paired t-test, n=2:4:51. *Bottom:* significant elevation of N/C ratio with prolonged and brief OGD (paired t-test). **(E)** Comparison of neuronal swelling caused by prolonged and brief OGD. Two-way ANOVA with Sidak’s post-test, F(1,188)=19.8, p<0.0001, n as D. **(F)** *Left:* Significantly larger elevation in N/C ratios with brief OGD (OGD+reperf) compared to prolonged OGD. Two-way ANOVA with Sidak post-test, F(1,188)=70, p<0.0001). *Right:* Cumulative distributions of N/C ratios of all neurons at baseline (-5’) and after prolonged and brief OGD (OGD+reperf) at 20’. Data represented as mean ± 95% CI (parametric data) or median ± IQR (non-parametric data). Scale bar: 50 µm.

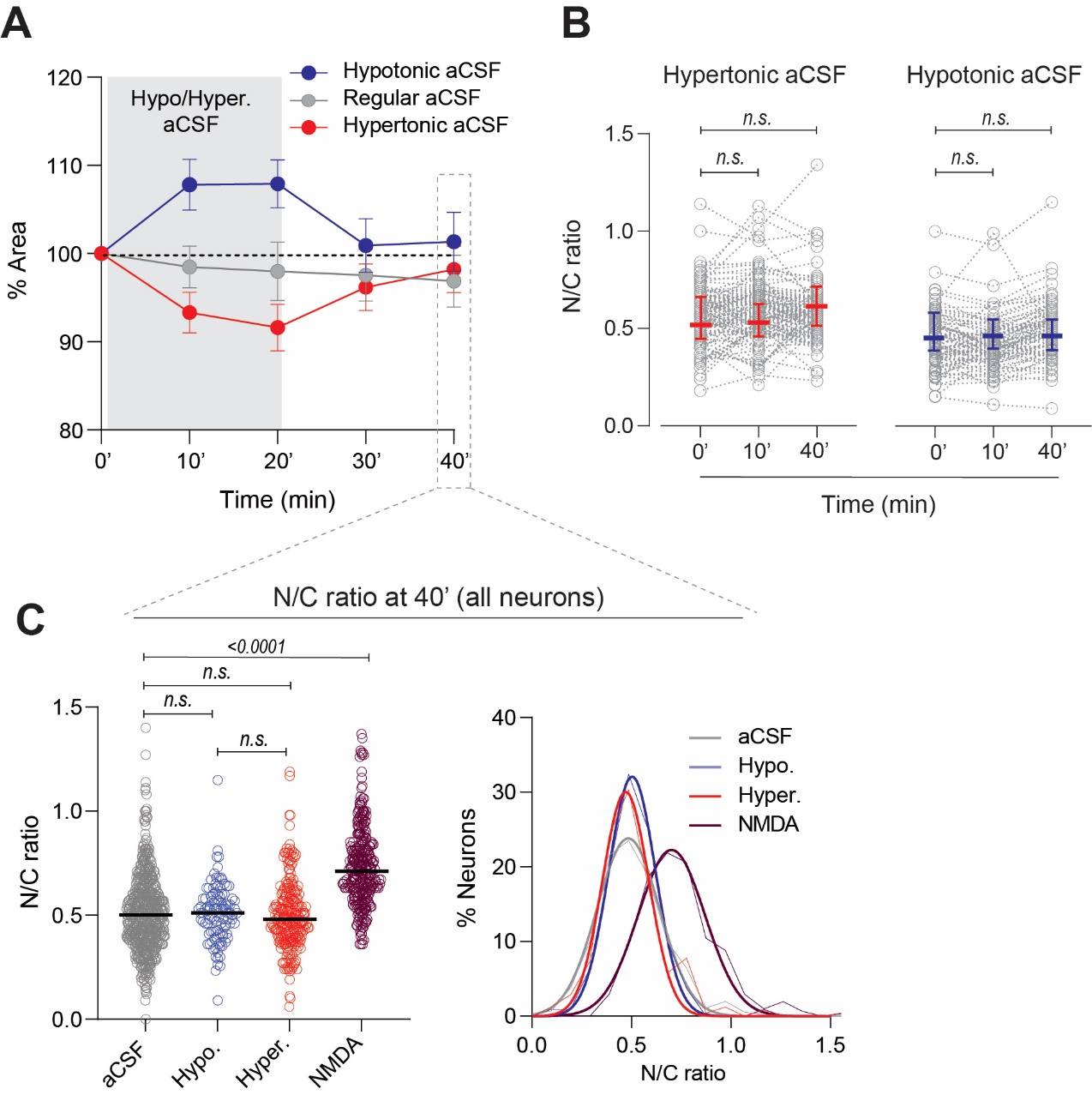

**Fig. S7. Neuronal osmotic swelling or shrinkage does not increase the N/C ratio.**

**(A)** Transient changes in neuronal areas induced by osmotic perturbations (blue: hypotonic -20 mOsm, red: hypertonic +40 mOsm). **(B)** No change in N/C ratios measured from matched neurons across time in hypo/hypertonic conditions. Friedman test with Dunn’s post-test; *left*: hypertonic, F(1.99, 213)=1.86, p=0.15; *right*: hypotonic, F(1.84, 145)=0.16, p=0.86; n= 3:7:85 (hypotonic), n= 3:7:112 (hypertonic). **(C)** *Left:* N/C ratios measured from all neurons at 40’ show no increase after hypotonic or hypertonic aCSF, unlike NMDA treatment. Kruskal Wallis test with Dunn’s post-test; n (total neurons), aCSF= 579, hypotonic= 124, hypertonic= 243, NMDA= 315). *Right*: N/C ratios from all neurons at 40’ show overlapping distributions in regular, hypertonic, and hypotonic aCSF conditions, compared to NMDA. Data represented as mean ± 95% CI (parametric data) or median ± IQR (non-parametric data).

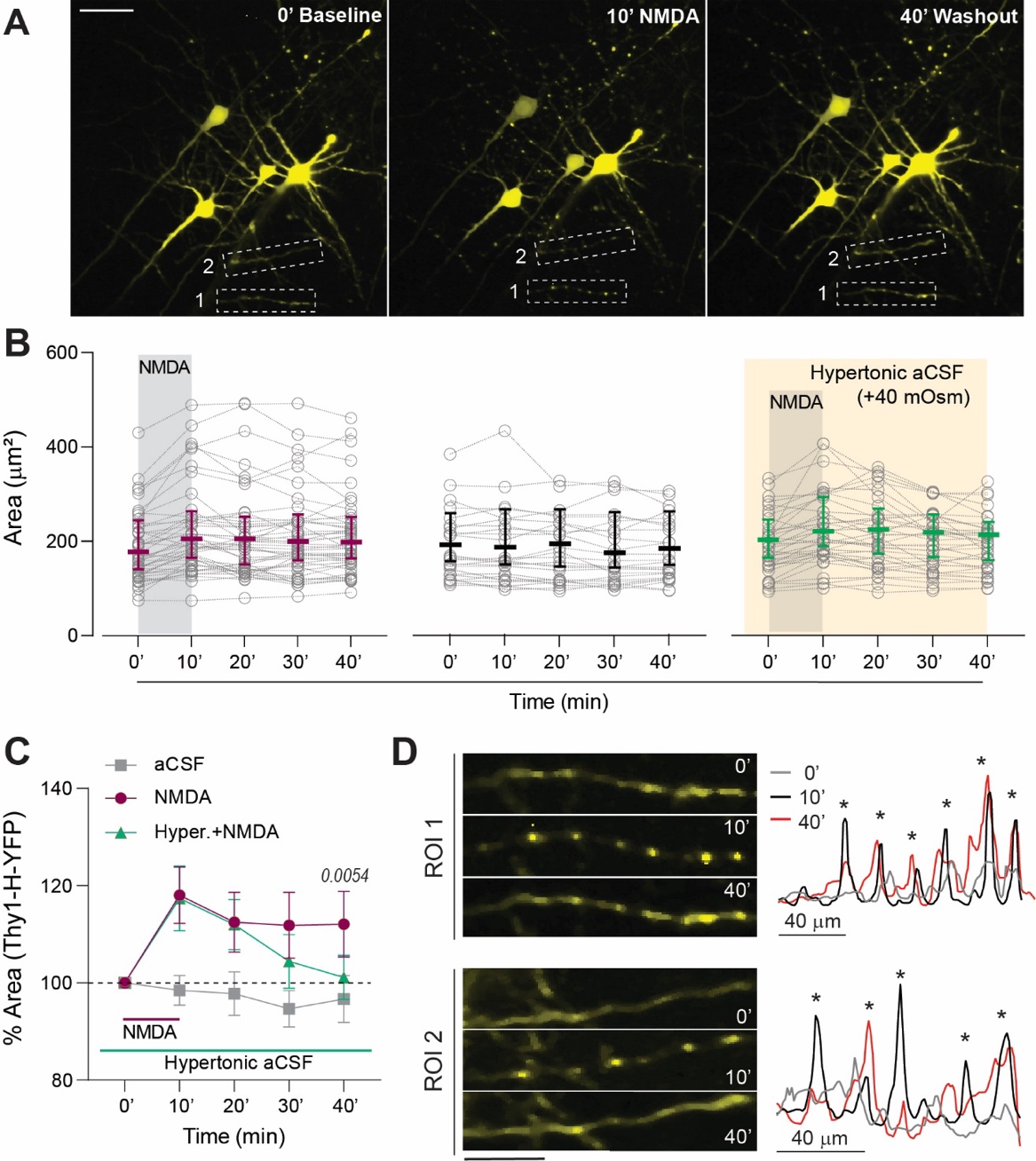

**Fig. S8. Hypertonic aCSF mitigates NMDA-induced neuronal edema in neonatal YFP-expressing neurons.**

**(A)** Sparse YFP-expressing neurons under the Thy1 promoter at 0’, after 10’ of NMDA perfusion, and during washout 40’ in hypertonic solution. ROI 1 and 2 show dendrites with semi-reversible varicose beading (see **D**), a morphological feature of excitotoxic injury. **(B)** Area changes in matched neurons over time in NMDA (magenta), aCSF alone (black), and NMDA + hypertonic aCSF conditions (green). n (mice: slices: neurons), NMDA = 3:6:46, aCSF = 3:4:25, NMDA + hypertonic aCSF =3:9:37. **(C)** Hypertonic aCSF significantly reduced NMDA-mediated neuronal swelling at 40’. Two-way ANOVA with Tukey’s post-test*,* Interaction: F (8, 525) = 2.99, p=0.0028; Time: F (4, 525) =7.6, p<0.0001; Treatment: F (2, 525) =28.9, p<0.0001. **(D)** *Left:* Magnified dendrites (ROI1 and 2 from A) showing varicose dendritic beading following NMDA treatment at 10’. *Right:* YFP signal intensities at linear ROIs drawn over neuronal dendrites show localized intensity peaks (*) at varicose beads at 10’ and 40’. Data represented as mean ± 95% CI (parametric data) or median ± IQR (non-parametric data). Scale bar: 50 µm (25 µm for **D**).

**Multimedia files**

**Video S1. Synchronous Ca^2+^ activity during NMDA perfusion. (Related to Fig. 1).**

Example of synchronous Ca^2+^ activity induced by NMDA application in a brain slice expressing GCaMP6s (green channel) and mRuby2 (red channel) under Synapsin-1 promoter. The video shows neurons at baseline (aCSF perfusion) followed by synchronous Ca^2+^ activity evoked by NMDA application. Data were acquired at 2.67 Hz. A median filter was applied (radius:2) for clarity. Playback speed: 30×. Scale bar (white): 50 μm.

**Video S2. NMDA application induced Ca^2+^ activity and nuclear translocation of GCaMP6s *in vivo.* (Related to Fig. 3).**

Example of transient Ca^2+^ activity induced by NMDA application followed by a delayed nuclear translocation of GCaMP6s during washout *in vivo*. The video shows GCaMP6s expressing neurons at baseline, Ca^2+^ activity evoked by NMDA application, followed by GCaMP6s translocation into neuronal nuclei during washout. Data was acquired from a neonatal Thy1-GCaMP6s pup (P11) at 2.67 Hz. A median filter was applied (radius:2) for clarity. Playback speed: 18×. Scale bar (white): 50 μm.

**Video S3. Ca^2+^ independent isosbestic excitation of GCaMP6s*.* (Related to Fig. S5).**

Imaging GCaMP6s expressing neurons following excitation at 920nm (left panel) and 810nm (right panel, two-photon isosbestic excitation) in an acute brain slice. The laser power (Bias and Pockels cell voltage) was consistent for both wavelengths. The video shows Ca^2+^ transients evoked by two electrical stimuli, clearly discernable at 920nm, yet unrecognizable at 810nm, demonstrating Ca^2+^ independent excitation of GCaMP6s. Data were acquired at 2.67 Hz. A median filter was applied (radius:2) for clarity. Playback speed: 34×. Scale bar (white): 50 μm.

**Video S4. Ca^2+^ transients evoked by repeated NMDA puffs*.* (Related to Fig. 4).**

Example of transient Ca^2+^ activity evoked by multiple NMDA puffs (1mM, 100ms, 3psi). The video indicates the exact number of puffs (stimulus #) associated with each Ca^2+^ transient. Data were acquired at 2.67 Hz. A median filter was applied (radius:2) for clarity. Playback speed: 24×. Scale bar (white): 50 μm.

**Video S5. Neuronal Ca^2+^ transients during 4-AP induced seizure-like activity*.* (Related to Fig. 4).**

Example of neuronal Ca^2+^ transients evoked by prolonged 4-AP perfusion (100µM, ~70 mins) inducing seizure-like activity in an acute brain slice. The video shows neuronal Ca^2+^ activity acquired at baseline and at multiple time points during 4-AP application. Data were acquired at 2.67 Hz (2 mins at each time point, then concatenated). A median filter was applied (radius:2) for clarity. Playback speed: 60×. Scale bar (white): 50 μm.

**Video S6. Anoxic depolarization (AD) induced by oxygen-glucose deprivation (OGD)*.* (Related to Fig. S6).**

Example of neuronal Ca^2+^ elevation associated with anoxic depolarization (AD) induced by oxygen-glucose deprivation (OGD, AD onset ~8.5 mins). Left: raw image stack. Right: ΔF/F_0_ stack. Data were acquired at 2.67 Hz. A median filter was applied (radius:2) for clarity. Playback speed: 24×. Scale bar (white): 50 μm.
